# Spatial representability of neuronal activity

**DOI:** 10.1101/2021.08.08.455535

**Authors:** D. Akhtiamov, A. G. Cohn, Y. Dabaghian

## Abstract

A common approach to interpreting spiking activity is based on identifying the firing fields—regions in physical or configuration spaces that elicit responses of neurons. Common examples include hippocampal place cells that fire at preferred locations in the navigated environment, head direction cells that fire at preferred orientations of the animal’s head, view cells that respond to preferred spots in the visual field, etc. In all these cases, firing fields were discovered empirically, by trial and error. We argue that the existence and a number of properties of the firing fields can be established theoretically, through topological analyses of the neuronal spiking activity.

## I. INTRODUCTION AND THE PHYSIOLOGICAL BACKGROUND

Physiological mechanisms underlying the brain’s ability to process spatial information are discovered by relating parameters of neuronal spiking with characteristics of the external world. In many cases, it is possible to link neuronal activity to geometric or topological aspects of a certain space—either physical or auxiliary. For example, a key insight into neuronal computations implemented by the mammalian hippocampus is due to O’Keefe and Dostrovsky’s discovery of a correlation between the firing rate of principal neurons in rodents’ hippocampi and the animals’ spatial location [1–3]. This discovery allowed interpreting these neurons’ spiking activity, henceforth called *place cells*, as representations of spatial domains—their respective *place fields* (Fig. 1A, [4]). ^1^ It then became possible to use place field layout in the navigated environment 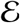— the *place field map* 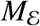— to decode the animal’s ongoing location [5–8], and even to interpret the place cells’ off-line activity during quiescent stages of behavior or in sleep [9–14], which define our current understanding of the hippocampus’ contribution to spatial awareness [15–18].

**FIG. 1:**
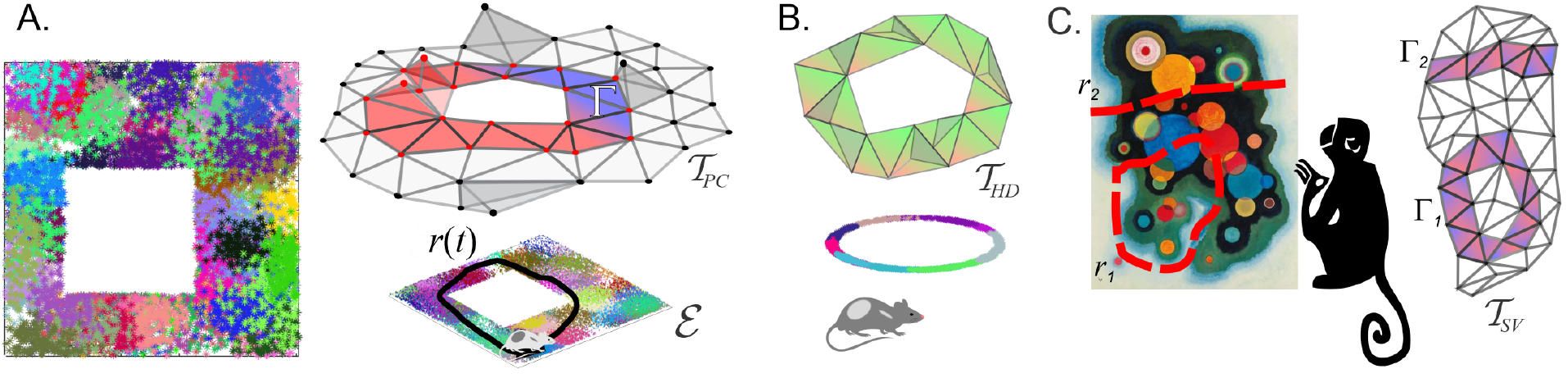
Spatial maps. (**A**) A simulated place field map of a small (1*m* × 1*m*) environment 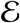, similar to the arenas used in typical electrophysiological experiments [66, 67]. Dots represent spikes produced by the individual cells (color-coded); their locations mark the rat’s position at the time of spiking. The pool of place cell coactivities is schematically represented by a coactivity complex 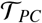 (top right). The navigated trajectory *r*(*t*) induces a sequence of activated simplexes—a simplicial path 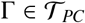. (**B**) The head direction cell combinations ignited during navigation induce a coactivity complex 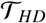 (top). The corresponding head direction fields cover a unit circle—the space of directions (bottom). (**C**) Spatial view cells activate when the primate gazes at their respective preferred domains in the visual field (left). The curves *r*_1_(*t*) and *r*_2_(*t*) traced by the monkey’s gaze induce simplicial paths Γ_1_ and Γ_2_ running through the corresponding coactivity complex 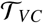 (right).

In the 90s, a similar line of arguments was applied to cells discovered in rat’s postsubiculum and in other parts of the brain [19–21], which fire at a particular orientation of the animal’s head. The angular domains where such *head direction cells* become active can be viewed as one-dimensional (I*D*) *head direction fields* in the circular space of planar directions, *S*^1^—in direct analogy with the hippocampal place fields in the navigated space (Fig. 1B). The corresponding *head direction map*, *M*_*S*1_, defines the order in which the head direction cells spike during the rat’s movements and the role of these cells in spatial orientation [20–22]. Recently, place cells and head directions cells were discovered in bats’ hippocampi; in contrast with rodents who navigate two-dimensional (2*D*) surfaces (see however [23–26]), bat’s voluminous place fields cover three-dimensional (3*D*) environments and their head direction fields cover 2*D* tori [27, 28].

The *spatial view cells*, discovered in the late 90s, activate when a primate is looking at their preferred spots in the environment (Fig. 1C), regardless of the head direction or location [29–31]. Correlating these cells’ spike timing with the positions of the *view fields* helped understanding mechanisms of storing and retrieving episodic memories, remembering object locations, etc. [32,33]. The principles of information processing in sensory and somatosensory cortices were also deciphered in terms of receptive fields—domains in sensory spaces, whose stimulation elicits in spiking responses of the corresponding neurons [34–39].

In all these cases, referencing an individual neuron’s activity to a particular domain in a suitable *representing space X* [40] is key for understanding its contribution and for reasoning about functions of neuronal ensembles in terms of the corresponding “maps”[16–18]. This raises a natural question: when is a “spatial” interpretation of neuronal activity at all possible, i.e., when there might exist a correspondence between the patterns of neuronal activity and regions in lowdimensional space?

## II. APPROACH

**A mathematical perspective** on this question is suggested by the simplicial topology framework [41, 42]. Specifically, if a combination of coactive cells, *c*_*i*0_, *c*_*i*1_, …, *c_ik_* is represented by an abstract *coactivity simplex* (for definitions see Sec. V)

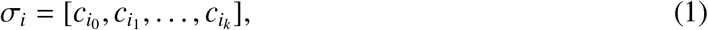

then the net pool of coactivities observed by the time *t* forms a simplicial complex

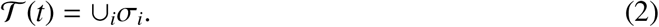

On the other hand [42–45], a similar construction can be carried out for a space *X* covered by a set of regions *ν_i_*,

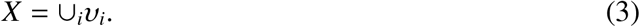

If each nonempty overlap between these regions,

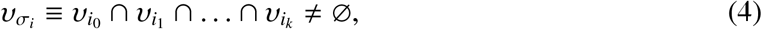

is formally represented by an abstract simplex,

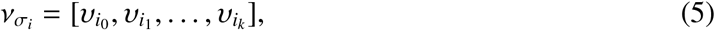

then the cover (3) generates another simplicial complex, known as its *Čech* or *nerve* complex

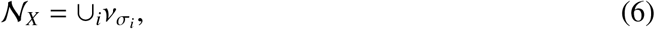

which is a spatial analogue of the coactivity complex (2). The idea is hence the following: if there is a correspondence between neurons’ spiking and spatial regions, then multi-cell coactivities can be viewed as representations of their firing fields’ overlaps [46–48]. Thus, the question whether a given pool of neuronal activity corresponds to a spatial map can be answered by verifying *representability* of the corresponding coactivity complex 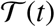, i.e., testing whether the latter has a structure of a nerve 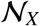 of some cover in a low-dimensional representing space *X*.

### Implementation

As it turns out, representable simplicial complexes exhibit several characteristic properties that distinguish them among generic simplicial complexes [49, 50]. Verifying these properties over biologically relevant 1*D*, 2*D* and 3*D* representing spaces is a tractable problem [49, 51], although exact algorithms for performing such a verification are not known—only in 1D are some methods available [53–57]. Nevertheless, there exist explicit criteria that allow limiting the dimensionality of the representing space *X* and eliminating manifestly non-representable complexes based on their homologies, combinatorics of simplexes and other intrinsic topological properties, which will be used below.

Specifically, according to the *Leray criterion*, a complex Σ representable in *D* dimensions should not contain non-contractible gaps, cavities or other topological defects in dimensionalities higher than (*D* − 1) [58]. Formally, it is required that the homological groups of Σ and hence its Betti numbers should vanish in these dimensions, *b*_*i*≥*D*_(Σ) = 0. Moreover, the Betti numbers of all the subcomplexes Σ_*x*_ of Σ, induced by a fraction *x* of its vertexes should also vanish, *b*_*i*≥*D*_(Σ_*x*_) = 0. In the case of coactivity complexes, such subcomplexes 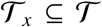 have a particularly transparent interpretation: they are the ones generated by *x*% of the active cells. According to the second criterion, the number of simplexes in all dimensions of Σ must obey *Eckhoff’s inequalities*— a set of combinatorial relationships discussed in [59–62] and listed in the Sec. V, where we also briefly detail the Leray criterion [49, 58, 62–64].

Previous topological studies of the coactivity data were motivated by the Alexandrov-Čech theorem [42–45], which posits that the homologies of the nerve complexes produced by the “good” covers (i.e., the ones with contractible overlaps (4), see [65]), should match the homologies of the underlying space *X*, 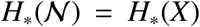, i.e., have the same number of pieces, holes, tunnels, etc. Specifically, this construction was applied to the place cell coactivity complexes, whose representability was presumed [46–48]. Persistent homology theory [68–73] was used to trace the dynamics of the Betti numbers 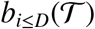 in physical dimensionalities *D* ≤ 2 [74–79] and *D* ≤ 3 [79, 80], to detect whether and when these numbers match the Betti numbers of the environment, 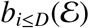, and how this dynamics depends on spiking parameters. It was demonstrated, e.g., that for a wide range of the firing rates and place field sizes referred to as the *Learning Region*, 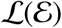, the low-dimensional Betti numbers of 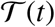 converge to their physical values after a certain period *T*_min_, neurobiologically interpreted as the minimal time required to “learn” the topology of the environment 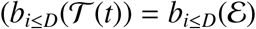, *t* ≥ *T*_min_ [48]).

Moreover, it became possible to asses the contribution of various physiological parameters— from brain waves to synapses—to producing and sustaining the topological shape of 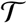 [74–80]. In addition, the coactivity complexes were used for contextualizing the ongoing spiking activity and linking its structure to the animal’s behavior. For example, it was shown that a trajectory *γ*(*t*) tracing through a sequence of firing domains *ν_σ_i__*, produces a “simplicial path” Γ—a succession of active simplexes that captures the shape of *γ*(*t*) and allows interpreting the animal’s active behavior [5–8] and its “off-line” memory explorations [9–14, 20, 32] (Fig. 1).

Together, these arguments suggest that experimentally discovered representing spaces and firing fields serve as explicit models of the cognitive maps emerging from neuronal activity—a perspective that is currently widely accepted in neuroscience. However, this view requires verification, since the empirically identified firing fields may be contextual offshoots or projections from some higher-dimensional constructs—in the words of H. Eichenbaum, “*hippocampal representations are maps of cognition, not maps of physical space*” [81]. The way of addressing this question is straightforward: if the spiking activity is intrinsically spatial, i.e., if neurons represent spatial domains, then the coactivity complexes generated by the corresponding neuronal ensembles should be representable—an explicit property that can be confirmed or refuted using Leray, Eckhoff and other criteria. In the following, we apply these criteria to several types of neuronal activity, both simulated and experimentally recorded, and discuss the results.

## III. RESULTS

### Simplicial topology approach

The conventional theory of representability addresses properties of “static” simplicial complexes [49–65]. In contrast, the coactivity complexes are dynamic structures that can be viewed as time-ordered agglomerates of simplexes, restructuring at the moments *t*_1_ < *t*_2_ < *t*_3_…,

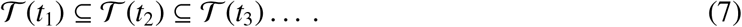

The exact organization of each complex in the sequence (7) depends on the specifics of the underlying spiking activity, e.g., the initial state of the network, its subsequent dynamics, spiking mechanisms and so forth (in case of the place fields, think of the starting point of navigation, shape of the trajectory, speed, etc.). Thus, verifying representability of these complexes requires testing whether Eckhoff, Leray and other criteria are valid at each moment *t*.

We constructed coactivity complexes by simulating the rat’s navigation through a planar environment 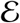 commonly used in electrophysiological experiments (Fig. 1A, see also [66, 67]). The neuronal spikings in this case are generated as responses to the rat’s appearances within preconstructed, convex firing domains, e.g., stepping into randomly scattered place fields or facing towards head direction fields centered around randomly chosen preferred angles (see Sec. V and Methods in [48, 74, 82]). While the resulting nerve complexes (6) are 2*D*-representable by design, we inquired whether the corresponding coactivity complexes are also representable, i.e., whether the activity of individual neurons intrinsically represents regions and whether connectivity between these regions is similar to the connectivity between the underlying auxiliary firing fields.

Simulations show that *persistent Leray dimensionality* 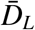 (above which the spurious loops in 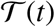 vanish, 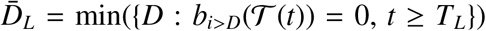, see also [83]) eventually settles at 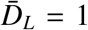 for most complexes, implying that neuronal activity defines a proper planar map. However, this mapping requires time—a *Leray period T_L_*—which, for the maps populating the learning region 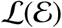, is typically similar to the learning time *T*_min_ (Fig. 2A,B).

**FIG. 2:**
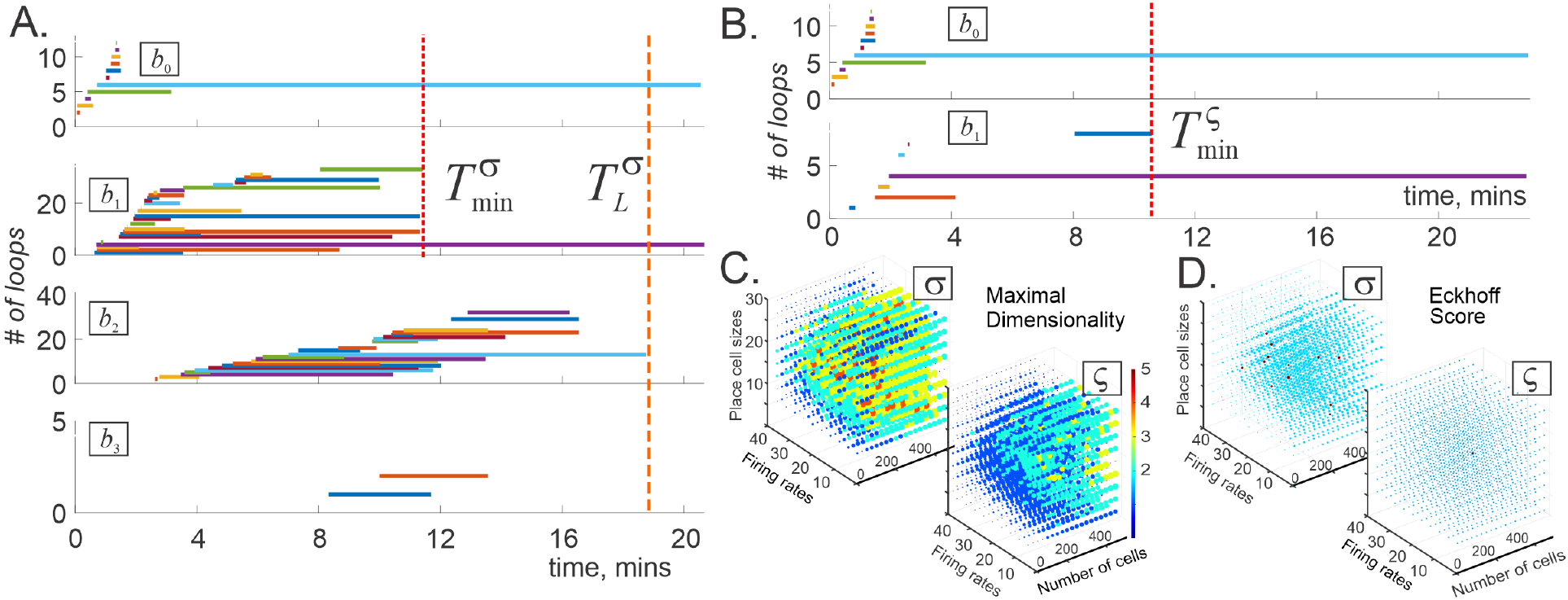
Persistent Leray dimension. (**A**). The Leray dimensionality of the coincidence-detector complex 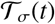 constructed for an ensemble of *N_c_* = 300 place cells can rise to 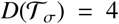 (here the mean maximal firing rate is *f* = 12 Hz, mean place field size *s* = 22 cm; environment 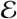 same as on Fig. 1A). In about 17 minutes—the corresponding Leray period *T_L_*— the dimensionality drops to 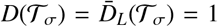, after which the spiking patterns can be intrinsically interpreted in terms of planar firing fields. Note that the Leray period in this case is longer than the minimal learning time evaluated based on the lower-dimensional Betti numbers 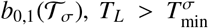. Shown are all the non-zero Betti numbers of 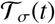. (**B**). Timelines of the topological loops in a spike-integrating coactivity complex, evaluated for the same cell population in the same environment yields the persistent Leray dimensionality 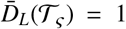 from the onset. The disappearance of spurious 0*D* loops in about 11 minutes marks the end of the learning period 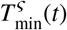. Note that the number of spurious loops in 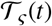 is significantly lower than in 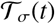. **C**. Maximal dimensionality of the topological loops in 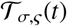. (**D**). The Eckhoff conditions are satisfied for nearly all coincidence-detecting complexes 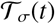 (left panel, occasional exceptions are shown by red dots) and for all spike integrating complexes 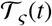 (right panel).

Whether a particular value of *T_L_* is shorter or longer than the corresponding *T*_min_ depends on how exactly the coactivity complex is constructed, e.g., whether the simplexes (1) correspond to simultaneously igniting cells groups or assembled from lower-order combinations over an extended period *ϖ* [84]. Physiologically, the former corresponds to the case when spiking outputs are processed by “coincidence-detector” neurons in the downstream networks and the latter to the case when lower-order coactivities are collected over a certain “spike integration window” *ϖ*—longer than the simultaneity detection timescale *w* [85–87]. Different readout neurons or networks may have different integration periods; to simplify the model, we started by extending the parameter ϖ to the entire navigation period for all cells and cell groups.

The lowest order of coactivity involves spiking cell pairs [88], which together define a coactivity graph 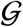 [89, 90]. The cliques *ς* of this graph produce a *clique coactivity complex* 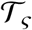 that generalizes the *simplicial coactivity complex* 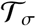, built from simultaneously detected simplexes [75–80]. As it turns out, the “coincidence detecting” and the “spike integrating” complexes have different topological dynamics: the former are more likely to start off with a higher Leray dimensionality, *D_L_* ≥ 3, that then reduces to 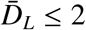 (Fig. 2A), whereas the latter tend to be more stable, lower-dimensional and have shorter Leray and learning times (Fig. 2B).

To test the induced subcomplexes of each 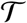, we selected random subcollections of cells containing *x* = 50%, *x* = 33%, *x* = 25% and *x* = 20% of the original neuronal ensemble, and found that if the original complex 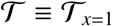 is representable, then its subcomplexes, 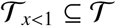, typically require less time to pass the Leray criterion, 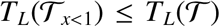, and that for *x* > 50% the Leray times saturate, 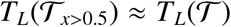. Thus, the Leray time of the full complex, 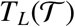, can be used as a general estimate of the timescale required to establish representability.

To control the sizes of the coactivity complexes, we used only those periods of each neuron’s activity when it fired at least *m* spikes per coactivity window (*w* ≈ 1/4 secs; for justification of this value see [74, 92]). Additionally, we used only those groups of coactive cells in which pairwise coactivity exceeded a threshold *μ* (Sec. V). Biologically, these selections correspond to using only the most robustly firing cells and cell assemblies for constructing the coactivity complexes [77]. The results demonstrate that majority of the coactivity complexes 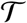 computed for smallest possible *m* and *μ* exhibit low *persistent Leray dimensionality*, 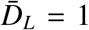, which points at 2*D* representability of the underlying neuronal activity, with the Leray times *T_L_* similar to the corresponding learning times *T*_min_ (Fig. 2C). We also found that Eckhoff inequalities are typically satisfied throughout the navigation period, i.e., that the Eckhoff criterion does not significantly limit the scope of representable spiking in this case (Fig. 2D).

### 2. Region Connection Calculus (RCC)

An independent perspective on spatial representability is provided by Qualitative Space Representation approach (QSR, [93, 94]), which sheds a new light on the dynamics of neuronal maps. From QSR’s perspective, a population of cells 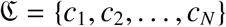 may represent a set of abstract, or *formal* spatial regions *R* = {*r*_1_, *r*_2_,…, *r_N_*}, if the relationships between them, as defined by the cells’ coactivity, can be consistently actualized in a topological space *X* by a set of explicit regions, ϒ = {*ν*_1_, *ν*_2_,… *ν_N_*}.

Specifically, regions *r_i_* and *r_j_* encoded by the cells *c_i_* and *cj* can be:

1. disconnected, DR(*r_i_, r_j_*), if *c_i_* and *c_j_* never cofire;
2. equal, EQ(*r_i_, r_j_*), if *c_i_* and *c_j_* are always active and inactive together;
3. proper part of one another, if *c_j_* is active whenever *c_i_* is, PP(*r_i_, r_j_*), or vice versa, PPi(*r_i_, r_j_*);
4. partially overlapping, PO(*r_i_, r_j_*), if *c_i_* and *c_j_* are sometimes (but not always) coactive.

These five relations fully capture mereological configurations of regions in a first-order logical calculus known as RCC5 (Fig. 3A, [95]). Using mereological, rather than topological, distinctions reflects softness of the firing fields’ boundaries: the probabilistic nature of neuronal spiking does not warrant determining whether two regions actually abut each other or not.

**FIG. 3:**
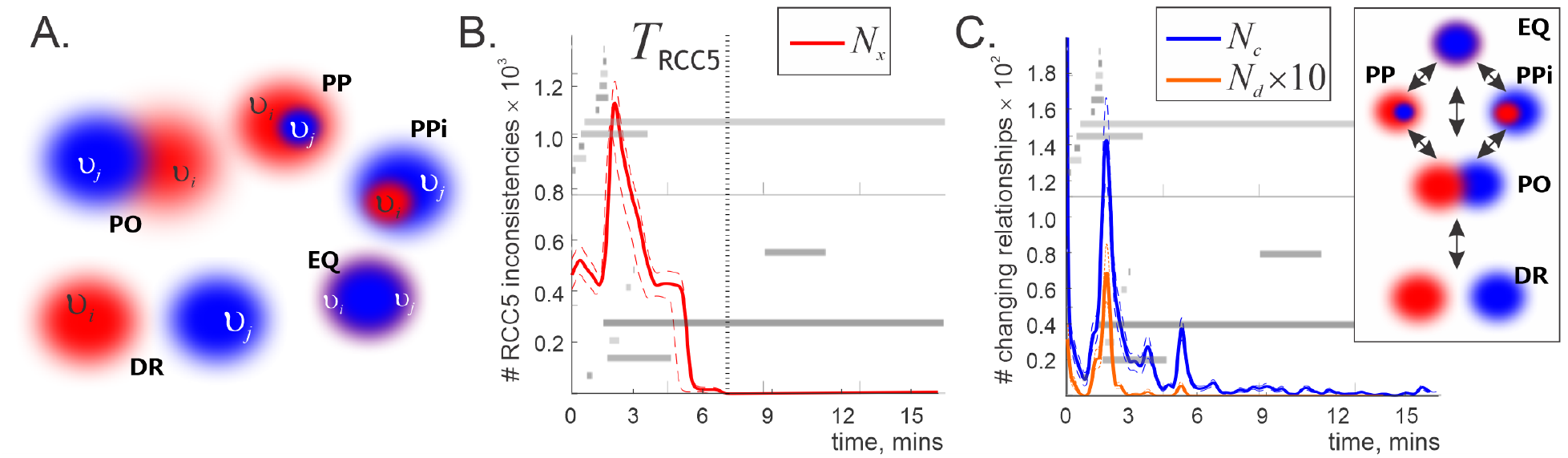
RCC5 analyses. **A**. Two regions with soft boundaries, e.g. two firing fields *ν_i_* and *ν_j_*, can overlap, PO(*ν_i_, ν_j_*), be proper parts of each other, PP(*ν_i_, ν_j_*) or PPi(*ν_i_, ν_j_*), be disconnected DR(*ν_i_, ν_j_*) or coincide EQ(*ν_i_, ν_j_*). **B**. Number *N_x_*(*t*) of inconsistent triples of RCC5 relationships appearing in the 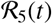 relational framework constructed for the same neuronal ensemble as illustrated in Fig. 2. The barcode diagram for the corresponding integrating coactivity complex (Fig. 2B) is shown in the background, to illustrate the correspondence between the RCC5 and the homological dynamics. *T*_RCC5_ (dotted line) marks the time when inconsistencies in the 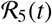 schema disappear. Results averaged over 10 repetitions, error margins shown by the dashed lines. **C**. The net number of changes of RCC5-relationships between two subsequent moments of time, *N_c_*(*t*), shown by the blue line, and the number *N_d_*(*t*) of changes that violate the RCC5 continuity order (top right panel), shown by the orange line. For better illustration, *N_d_*(*t*) is scaled up by a factor of 10. Initially, discontinuous events are frequent but shortly before *T*_RCC5_ they disappear entirely, leaving the stage to qualitatively continuous sequences. The same barcodes are added in the background, error margins shown by dashed lines.

A key property of a RCC5-framework defined by spiking neurons—a 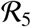 schema— is its internal consistency [40]. It may turn out, e.g., that some pairs of cells encode relationships that are impossible to reconcile, e.g., PP(*r_i_, r_j_*), DR(*r_j_, r_k_*) and PO(*r_i_, r_k_*). Indeed, if an actual region *ν_i_* is contained in *ν_j_* then it cannot possibly overlap with a region *ν_k_* that is disconnected from *ν_j_*. Correspondingly, the neuronal activity that produces such inconsistencies (for the full list see Table I in Sec. V) is not representable—not even interpretable in spatial terms. On the other hand, it can be shown that if all triples of relationships are consistent, then 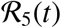 does possess a spatial model, i.e., there exists a set of regions *ν_i_* (with no prespecified properties such as convexity, connectivity or dimensionality) that relate to each other as the *r_i_*s relate in 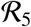 [93–98].

**TABLE I:**
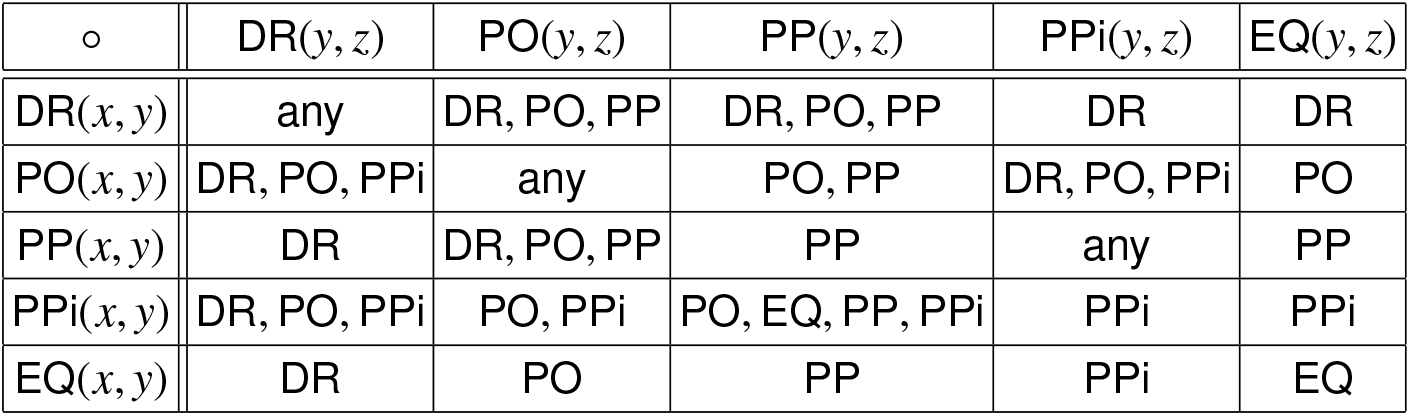
RCC5 **compositions**. Given three regions, *x, y* and *z*, and two relationships *R*_1_(*x, y*) and R_2_(*y, z*), the relationship R_3_(*x, z*) is not arbitrary. A map is consistent, if every triple of relationships is RCC5–consistent.

To verify whether spiking activity is representable in this QSR sense, we constructed an inflating 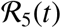-schema (an RCC5-framework growing as spiking data accumulates, similar to (2)) for each neuronal ensemble and counted the inconsistent triples of relationships at each moment *t*. The results show that all 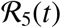-schemas start off with numerous inconsistencies, which tend to disappear after a certain period *T*_RCC5_ that is typically smaller than the Leray time *T_L_* (Fig. 3B,C).

The net dynamics of RCC5 relationships is illustrated on Fig. 3C. Note that some of these changes may be attributed to the regions’ continuous reshapings or displacements, e.g., two over-lapping regions may become disconnected, PO(*r_i_, r_j_*) → DR(*r_i_, r_j_*), is *r_i_* moves away from *r_j_*, or *r_i_* may move into *r_j_*, inducing PO(*r_i_, r_j_*) → PP(*r_i_, r_j_*). In contrast, a jump from a disconnect to a containment, without an intermediate partial overlap, e.g., DR(*r_i_, r_j_*) → PP(*r_i_, r_j_*) rather than DR(*r_i_, r_j_*) →PO(*r_i_, r_j_*) →PP(*r_i_, r_j_*), would be a discontinuous, abrupt change. As shown on Fig. 3C, discontinuous transitions are common at the initial stages of navigation, but shortly before *T*_RCC5_ they disappear, indicating that the relationships between regions encoded within a sufficiently well-developed 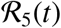 schema evolve in a continuous manner.

These outcomes not only provide an alternative lower-bound estimate for the time required to accumulate data for producing low-dimensional spatial representations, but also help understanding the nature of processes taking place prior to Leray time. In particular, the exuberant initial dynamics, homologically manifested through an incipient outburst of spurious loops in the coactivity complexes (Figs. 2A,B, *t* < *T*_min_), cannot be interpreted as a mere “settling” of topological fluctuations in the cognitive map—according to Fig. 3B, the RCC5-schema does not form a coherent topological stratum for 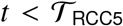. Rather, the initial disorderly period should be viewed as the time of transition from a nonspatial to a spatial phase, followed by spatial dynamics (for 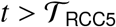) that involves, *inter alia*, dimensionality reduction and other restructurings (Fig. 3C).

### 3. Current summary

Taken together, these results show that even in the simplest “reactive” model, in which neuronal firings are simulated as responses to regular domains covering a compact space, the low-dimensional representability is not an inherent, but an emergent property. In particular, RCC5-analyses suggest that spatial interpretation of neuronal spiking becomes possible after a finite period. During the times that exceed both *T*_RCC5_ and *T_L_*, the spiking data can be interpreted in terms of firing fields in a space *X* of dimensionality higher than the persistent Leray dimensionality of the corresponding coactivity complex, 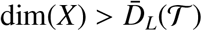.

Physiologically, this implies that the outputs of place cells, head direction cells, view cells, etc., may not be immediately interpretable by the downstream networks as representations of spatial regions—the information required for such inference appears only after a certain “evidence integration.” Correspondingly, the firing field maps constructed according to the standard experimental procedures [1, 19, 30] also cannot be considered as automatic “proxies” of cognitive maps: such interpretations are appropriate only after the representability of the corresponding coactivity complexes is established. Another principal conclusion is that representability of the spiking activity depends not only on the spiking outputs, but also on how the information carried by these spikes is detected and processed. In particular, spikes integrated over extended periods are likelier to permit a consistent firing field interpretation than spikes counted via coactivity detection. On the technical side, these results imply that an accurate description of the firing fields’ plasticity should include possible dimensionality changes [100–102].

### 4. Multiply connected place fields

A key simplification used in the simulations described above is that firing fields were modeled as convex regions. While this assumption is valid in some cases [99], multiply connected firing fields are also commonly observed (Fig. 4A, [103, 104]). From our current perspective, the issue is that multiple connectivity of the cover elements (3) may increase the Leray dimensionality of the corresponding nerve complex [49, 105] and thus bring additional ambiguity into the analyses. Identification of the firing fields’ connectivity from the spiking data is an elaborate task that requires tedious analyses of the spike trains produced by individual cells or cell groups over periods comparable to the Leray and the learning times [83, 106]. To circumvent these difficulties, we reasoned as follows.

**FIG. 4:**
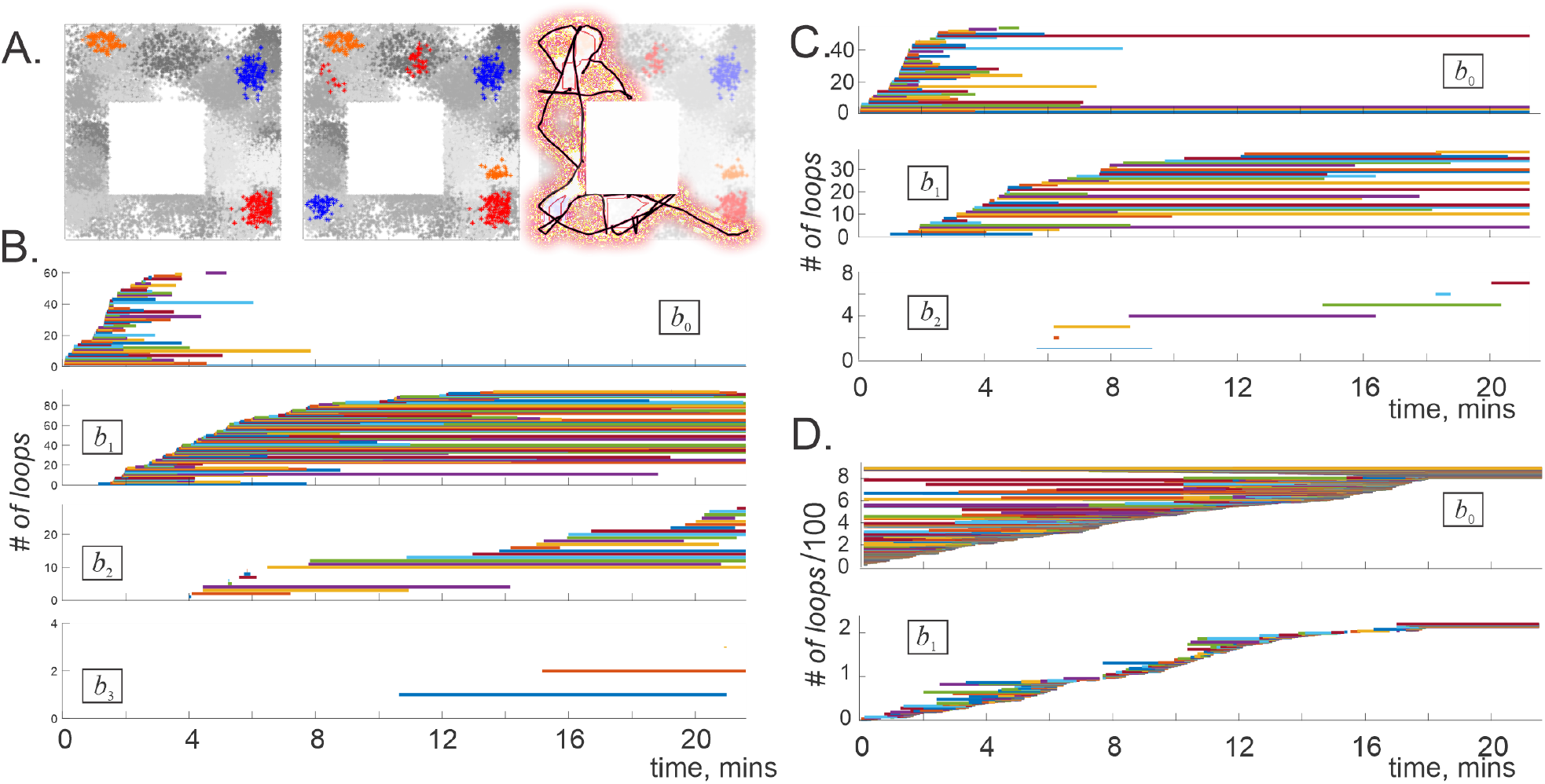
Topological dynamics in maps with multiple firing fields. (**A**). Left panel shows three examples of convex place fields used to obtain the results illustrated in Fig. 2. Allowing a cell to spike in several (2 − 3) locations produces multiply connected place fields (middle panel; clusters of dots of a given color correspond to spikes produced by a single simulated neuron). Right panel shows a *ϖ* = 50 second long fragment of the trajectory *ϓ_ϖ_* covering a segment *χ_ϖ_* of the environment (reddened area). (**B**). The Leray dimensionality of the detector-complex evaluated for the same place cell population as in Fig. 2A, with the exception of 30% of multiply connected place fields (2 − 3 components each) can reach 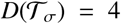. (**C**). In a clique coactivity complex, the spurious loops in dimensions *D* = 2 and lower may persist indefinitely, implying either that the firing fields are 3D-representable *or* that they may be multiply connected. Note that the number of spurious loops in both 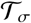 and in 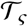 is higher than in the case with convex firing fields (Fig. 2A,B). (**D**). The persistence bars computed for the flickering complex 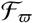 with spike integration window *ϖ* = 1 minute, indicate stable mean Leray dimensionality 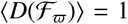, implying that the local charts *χ_ϖ_* are planar and hence that the firing fields are two-dimensional.

Suppose that the spiking activity used to produce a coactivity complex 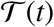 is generated as a moving agent (animal’s body, its head, its gaze) follows a trajectory *γ*(*t*) over a space *X*, covered with stable firing fields *ν_i_*, *i* = 1,…, *N*. Consider a navigation period *ϖ* that spans over a smaller segment of this trajectory, *γ_ϖ_* = {*γ*(*t*) : *t* ∈ *ϖ*}. If *ϖ* is sufficiently short, then one would expect *γ_ϖ_* to cross at most one component of a typical firing field *ν_i_*, even if 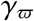 meets more than one component of a multiply connected *ν_i_*, this property may not manifest itself in the resulting spike trains, i.e., *ν_i_* should be *effectively* simply connected (Fig. 4A). Correspondingly, the Leray dimensionality of the coactivity complex acquired during that period should reflect the dimensionality of a small underlying fragment of *X*—a *local chart χ_ϖ_*—that contains *γ_ϖ_* (topologically, *χ_ϖ_*(*γ*) ≅ {∪_*j*_*ν*|*γ_ϖ_* ∩ *ν_j_* ≠ ⊘}). The dimensionality of *χ_ϖ_* can then be ascribed to all the contributing *ν_j_*s, dim(*ν_j_*) = dim(*χ_ϖ_*).

Further, if the *ϖ*-period is allowed to shift in time, then the segment *γ_ϖ_* will also slide along the trajectory *γ*(*t*); the spikes fired within each *t*-centered window, *ϖ* = [*t* – *ϖ*/2,*t* + *ϖ*/2], will then produce a *ϖ_t_*-specific *flickering coactivity complex* 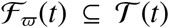, whose topological properties may change with time [78, 107, 108]. Since 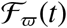 contains a finite number of elements, it will reconfigure at discrete moments, *t*_1_, *t*_2_,…, and remain unchanged in-between, 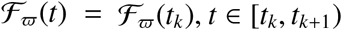, *t* ∈ [*t_k_*, *t*_*k*+1_). If a given instantaneous configuration 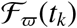 is representable, then its vertexes correspond to the regions comprising the local chart *χ_ϖ_*(*t_k_*), with dimensionality 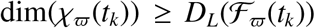. If two such complexes overlap, 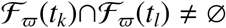 (i.e., their vertex sets overlap), then their respective charts also overlap *χ_ϖ_*(*t_k_*) ∩ *χ_ϖ_*(*t*_*k*+1_) ≠ ⊘, which allows relating their topological properties, including properties of the representing regions.

Clearly, the outcome may depend on how each 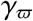 is embedded into *X*, the spiking parameters, etc. Moreover, since the Leray dimensionality of the instantaneous complexes can change, so can the dimensionalities of the corresponding local charts: 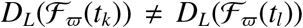 may entail dim(*χ_ϖ_*(*t_k_*)) ≠ dim(*χ_ϖ_*(*t_l_*)). This may seem as a contradiction since the representing space is naturally assumed to be a topological manifold, i.e., all of its local charts, arbitrarily selected, should have the same dimensionality dim(*χ_ϖ_*(*t*)) = dim(*X*) = *D*. On the other hand, the deviations of the local dimensionality estimates from a fixed *D* can be viewed as mere fluctuations caused by occasional contribution of multiply connected firing fields or by other noise sources, e.g., by stochasticity of neuronal spiking [109]. One can hence attempt to discover the true dimensionality of *X* by evaluating the mean Leray dimensionality of the instantaneous complexes,

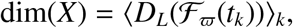

which physiologically alludes to learning the physical structure of the underlying space from the recurrent information.

Numerical verification of the viability of the proposed approaches can be achieved by simulating multiply connected firing fields and computing homological dynamics of the resulting coactivity complexes. To that end, we randomly added 2 − 3 additional convex components to ~ 30% of the place fields (Fig. 4A) and simulated the topological dynamics of the corresponding complexes.

The results show that multiple connectivity of the firing fields does indeed increase Leray dimensionality in both the detector and the integrator complexes, 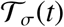 and 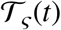. Moreover, in contrast with the complexes generated off the maps with convex fields, the maps with multiply connected fields tend to produce persistent higher-dimensional loops, notably in the coactivity detecting complexes 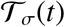 (compare Fig 2A and Fig. 4B). In the spike integrating clique complex 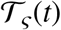, the Leray dimensionality remains low and may in some cases retain the physical value 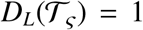, although topological loops in dimensions *D* = 2 and even higher may also appear (Fig. 4C). Thus, multiple firing field connectivity significantly increases the number of spurious 1*D* holes (by 200 – 300%), precluding both types of complexes from assuming the physically expected topological shapes.

Tighter dimensionality estimates can be produced by using shorter spike integration windows *ϖ* ≲ *T*_min_ and constructing flickering coactivity complexes 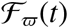 from pairwise coactivities detected over *ϖ*-periods shifting by discrete steps Δ*ϖ* and yielding an array of windows *ϖ*_1_, *ϖ*_2_, *ϖ*_3_,… centered at *t_k_ = ϖ*/2 + (*k* − 1)Δ*ϖ*. The specific *ϖ*-values were chosen comparable to the characteristic time required by the rat to run through a small segment of the environment: *ϖ* ≈ 25 − 65 secs for the arena shown on Figs. 1A and 4A. The Betti numbers for this case were evaluated using zigzag homology theory—a generalization of the persistent homology theory that applies to complexes that can not only grow, but also shrink, break apart, fuse back again, etc. [110, 111]. In particular, this approach allows studying how the topological fluctuations in 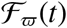 affect its Leray dimensionality 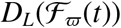 from moment to moment.

Typical results illustrated on Fig. 4D show that there appears a large number of spurious 0*D* loops—disconnected pieces—with lifetimes nearly exponentially distributed about the learning periods 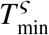, which suggests that fragments of 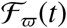 appear and disappear at random over such periods. The transient 1*D* loops also form and decay at *ϖ*-timescale. However, the most important outcome is that the topological dynamics in dimensions *D* > 1 trivializes—the higher dimensional loops in 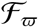 occur very rarely, if ever. These properties are qualitatively unaffected by varying the discretization step Δ*ϖ* (*ϖ*/20 ≲ Δ*ϖ* ≲ *ϖ*/10) or changing the window width *ϖ*, i.e., the estimates of the mean Leray dimensionality 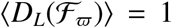 are stable and reveal physical planarity of the representing space.

Verification of the RCC5-consistency of the spiking data produces the same qualitative results as in the case with simply connected firing domains: the 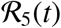-schemas become consistent after a learning period 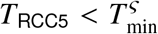, upon which neuronal activity becomes spatially interpretable, and, by the Leray and Eckhoff arguments, representable in dimensions 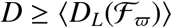.

### 5. Electrophysiological data

We applied the analyses described above to spiking activity recorded in the hippocampus (CA1 area) of rats navigating a linear environment shown on Fig. 5A (for more data description and experimental specifications see [112]). A typical running session, during which the animal performed 45 – 70 laps between the tips of the track, provided *N_c_* ≲ 25 simultaneously recorded neurons, allowing to construct small coactivity complexes that quickly become RCC5-consistent, comply with the Eckhoff conditions, and exhibit persistent Leray dimensionality, 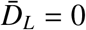, with typical persistent Leray time *T_L_* ≈ 10 mins (Fig.5B).

**FIG. 5:**
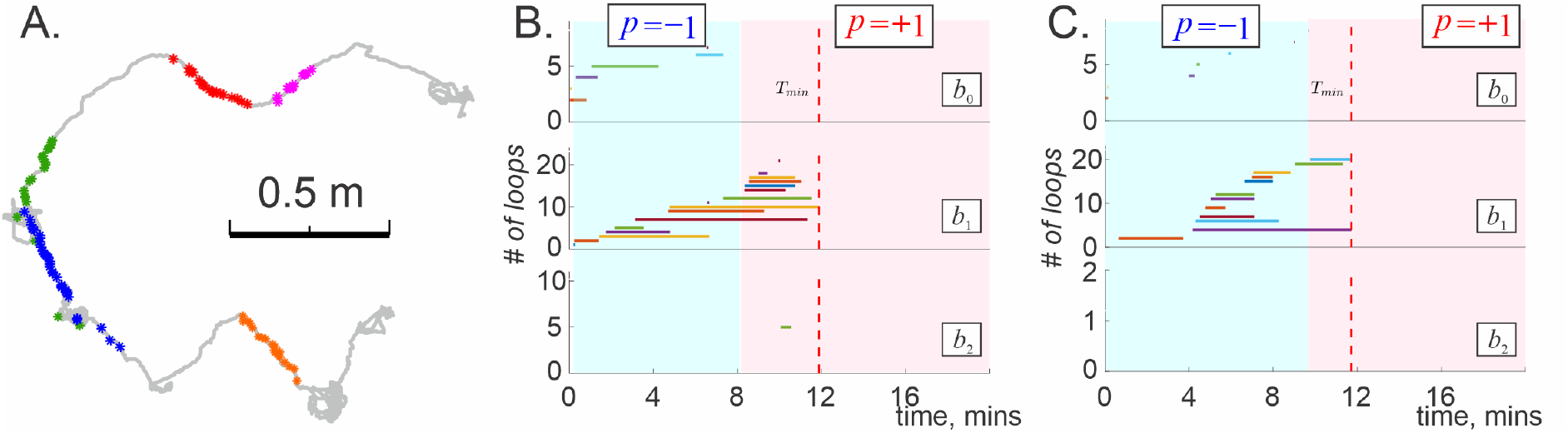
Multiple firing fields. (**A**) Spikes produced by five place cells (dots of different color) recorded in hippocampal CA1 area of a rat navigating a linear track (speed *ν* ≥ 3 cm/sec). The underlying gray line shows a fragment of the rat’s trajectory (for more details see [112]). (**B**) Spurious topological loops in the corresponding coactivity complex disappear in *T_L_* ≈ 12 minutes, revealing persistent Leray dimensionalities 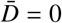. The blue background highlights the period during which the coactivity complex computed using only cells with convex place fields is not 1*D* representable (*p* = −1). The transition to *p*(*t*) = +1, marking the onset of 1*D* representability occurs at a time close to *T_L_*. (**C**) Topological dynamics of the coactivity complex constructed using the data recorded during the outbound moves only shows qualitatively similar behavior.

The vanishing 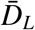 indicates that a linear track illustrated in Fig. 5A is contractible and implies 1*D*-representability. The latter can also be tested independently via RCC5 analyses, which in this case allows identifying the track’s linear structure [113].

Since some of the hippocampal place fields are multiply connected, we also applied sliding window analyses, adjusting the spike integration period *ϖ_i_* to match the duration of the animal’s *i*^th^ run from one end of the track to the other (typically 2 ≤ *ϖ_i_* ≤ 10 secs). Computations reveal that the resulting complexes 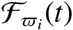 exhibit the same mean Leray dimensionality 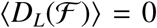, which is consistent with the persistent 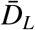 estimates above. Combining these results produces convergent evidence that in this case the hippocampus does indeed map out a 1*D* spatial domain (rather than 2D, see [112]).

The latter conclusion can, in fact, be verified by yet another representability test, which applies only to 1*D* cases and presumes firing field convexity. The Golumbic-Fishburn (GF) algorithm [53–57] is based on computing a binary index *p*: the 1*D*-representable simplicial complexes Σ yield *p*(Σ) = +1, and the non-representable complexes produce *p*(Σ) = −1. For the inflating or flickering coactivity complexes this index becomes time-dependent, *p* = *p*(*t*), marking the evolution of 1*D* representability (Sec. V). Applying the GF-algorithm to the inflating coactivity complexes constructed for cells with convex place fields only, we found that 1*D* representability, *p*(*t*) = +1, appears in about *T_+_* ≈ 10 mins, close to the Leray time (Fig. 5B,C), demonstrating consistency with the previously obtained results.

Lastly, we addressed a particular property of the place cell’s spiking activity in linear environments—the place fields’ directionality: a given place cell may fire during the outbound, but not inbound directions, or vice versa [114]. We verified that the topological dynamics exhibited by the coactivity complexes built using only the outbound or only the inbound activity are very similar to the dynamics of the full (bidirectional) complex 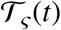 (Fig 5B,C), implying that place cell directionality does not necessarily compromise 1*D* representability of spiking activity.

## IV. DISCUSSION

Topological analyses of the spiking data allow testing whether a given type neuronal activity may arise from a “spatial map,” i.e., whether each neuron’s spiking marks a domain similar to a place field, a head direction field, a view field, etc., in a certain low-dimensional space. Thus far, establishing correspondences between neurons and firing fields was based on matching the spike trains with spatial domains empirically, through trial and error [1, 19, 30]. Here we attempt to address this question in a principled way, through intrinsic analyses of the spiking data, without presuming or referencing *ad hoc* constructions. A set of hands-off algorithms discussed above allows objective estimates for the dimensionality of a space needed to model the patterns of neuronal firing—a method that is unaffected by technical limitations, experimental ingenuity or complexity (e.g., nonlinearity) of the required firing field arrangements.

To follow the dynamics of the coactivity complexes we extend the conventional approaches of representability theory into the temporal domain, obtaining several complementary time-dependent markers of representability. In particular, we use persistent homologies to extend Leray’s theory to the case of inflating simplicial complexes and zigzag homologies in the case of flickering simplicial complexes. The latter approach is especially valuable as it allows extracting stable topological information from spiking data that may be generated from the maps with multiply connected firing fields or encumbered by other inherent irregularities, in spirit with the general ideas of topological persistence [68–73]. It should also be mentioned that mathematical discussions of the persistent nerve theorem, alternative to ours and more formal, have began to appear [115, 116]; however at this point our studies are independent.

A principal observation suggested by our analyses is that representability is a dynamic, emergent property that characterizes current information supplied by the neuronal activity. Moreover, representability depends not only on the amount and the quality of the spiking data itself, but also on the mechanisms used for processing and interpreting this data. Both aspects affect the time required to establish the existence of a representing space and its dimensionality. An implication of this observation is that experimentally constructed firing field maps (place field maps, head direction maps, etc.) cannot be automatically regarded as direct models of cognitive representations of ambient spaces [15–21] or more general spatial frameworks [117]; correctness of such interpretations may require more nuanced considerations.

## Acknowledgments

The authors would like to thank Dr. M. Tancer for fruitful discussions and valuable feedback and to D. Morozov for providing computational software. The work was supported by the European Research Council (ERC) under the European Unions Horizon 2020 Research and Innovation program, grant 692854 (D.A.), by Alan Turing Institute Fellowship and EPSRC under grant EP/R031193/1 (A.C.) and by NSF grant 1901338 (Y.D.)

## V. METHODS

### A. Physiological parameters and constructions

- *Simulated trajectory r*(*t*) = (*x*(*t*), *y*(*t*)), used for generating coactivity complexes was obtained by modeling a rat’s non-preferential exploratory behavior—navigation without favoring of one segment of the environment 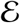 over another (Fig. 1A). The mean speed of about ~ 20 cm/sec was selected to match experimentally recorded speeds. The direction of the velocity *v*(*t*) = (*v_x_*(*t*), *v_y_*(*t*)) defines the “angular trajectory” *φ*(*t*) = arctan *v_y_*(*t*)/*v_x_*(*t*) that traverses the space of directions, *S*^1^, allowing to simulate head direction cell activity as the rat explores 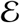 [48, 74, 82]. The simulated *navigation period, T* = 25 minutes, was selected to match the duration of a typical “running session” in electrophysiological experiments [99]. A shorter *spike integration window ϖ* ≪ *T* was used to limit the pool of spiking data for time-localized computations.
- *Poisson spiking rate* of a place cell *p* depends on the animal’s location *r*(*t*),

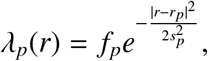

where *f_p_* is the cell’s maximal firing rate and *s_p_* defines the size of its place field [100]. A similar formula defines the firing rate of a head direction cell *h*, *λ_h_*(*φ*), as a function of the animal’s ongoing orientation *φ*, the cell’s preferred orientation angle *φ_h_*, its maximal rate *f_h_* and the size of its preferred angular domain *s_h_*. In all simulations the firing fields were stable, i.e., the parameters of *λ_c_* and *λ_h_* remained constant.
- *Neuronal ensembles* produce lognormal distributions of the maximal firing rate amplitudes, *f_c_*, and of the firing field sizes, *s_c_* [48, 118]. We tested about 17,000 different ensembles, in which the ensemble mean maximal rate *f* ranged between 4 and 40 Hz for the place cells and between 5 to 35 Hz for the head direction cells. The ensemble mean firing field sizes varied between 10 to 90 cm for the place fields and between 12°and 36°degrees for the angular fields. For all ensembles, the firing field centers were randomly scattered over their respective representing spaces.
- *Multiple Firing Fields* were generated by adding two or three randomly scattered auxiliary spiking centers *r*_*c*′_, *r*_*c*″_, etc.,

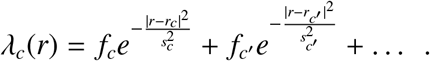 The maximal firing rates at the auxiliary locations are smaller than the rate at the main location, *f_c_* > *f*_*c*′_ > …, as suggested by the experiments [103, 104].
- *The activity vector* of a cell, *m_c_* = [*m*_*c*,1_,…, *m_c,n_*], is constructed by binning its spike trains into *w* = 1/4 seconds long “coactivity windows”[74, 92]. Each *m_c,k_* specifies how many spikes were fired by *c* into the *k^th^* time bin, *n* is defined by the duration of navigation, *n* = ⌊*T*/*w*⌋. High activity periods can be identified by selecting time bins in which the number of fired spikes exceeds an activity threshold *m*.
- *Coactivity*. Two cells, *c_i_* and *c_j_*, are *coactive* over a time period *T*, if the formal dot product of their activity vectors does not vanish, *m_ij_*(*T*) = *m_c_i__*(*T*) · *m_c_j__*(*T*) ≠ 0. The set of all pairwise coactivities forms the coactivity matrix *M*(*T*) = ||*m_ij_*(*T*)||. Highly coactive pairs of cells are the ones whose coactivity exceeds a threshold μ.

### B. Topological propaedeutics

#### Graphs

- *A graph G* is defined by its vertices, *V* = {*v*_1_, *v*_2_,…, *v_n_*}, and a set of edges *E* that link certain pairs of vertexes. A formal description of a graph is given by its connectivity matrix *C*(*G*), with the elements

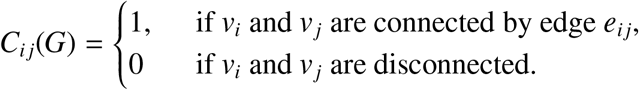
- *A coactivity Graph* 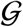 is built by establishing functional links between cells that exhibit high activity and coactivity (*m_c,k_* ≥ *m*, *m_ij_* ≥ *μ* see above) [89, 90].
- A *clique* of order *d*, *ς*^(*d*)^ in a graph is a fully interconnected subset of (*d* + 1) vertexes *v*_*i*_0__, *v*_*i*_1__,…, *v*_*i*_*d*__ (Fig. 6A).
- Given a graph *G*, its *complement graph* 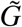 is produced by flipping 0s and 1s in the connectivity matrix *C*(*G*), i.e., joining the disconnected vertexes of *G* and removing the existing edges.
- A *comparability graph G*_◁_ represents an abstract relationship “◁”, if its vertexes *v_i_* represent elements of a set, and each link *e_ij_* represents a ◁-related pair, *v_i_* ◁*v_j_*.

**FIG. 6:**
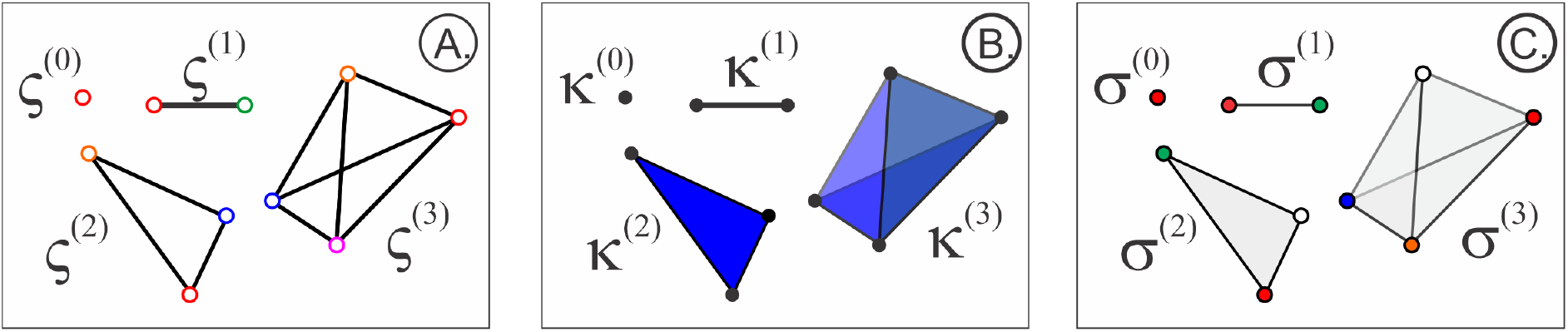
Cliques and simplexes. (**A**). Pairwise interlinked subsets of vertexes in graph *G* form its cliques. Shown is a vertex *ς*^(0)^ (0-clique), a link *ς*^(1)^ (1-clique), a three-vertex *ς*^(2)^ and a four-vertex *ς*^(3)^ cliques. (**B**). Geometric simplexes: a 0*D* dot (*κ*^(0)^), a 1*D* link (*κ*^(1)^), a 2D triangle (*κ*^(2)^) and a 3D tetrahedron (*κ*^(3)^). (**C**). The corresponding abstract simplexes: *σ*^(0)^ (vertexes), *σ*^(1)^ (pairs of vertexes), *σ*^(2)^ (triples) and *σ*^(3)^ (quadruples).

#### Simplicial complexes

- *Geometric simplexes* are points (0-simplexes, *κ*^(0)^), line segments (1-simplexes, *κ*^(1)^), triangles (2-simplexes, *κ*^(2)^), tetrahedra (3D-simplexes, *κ*^(3)^), as well as their *d* >3-dimensional generalizations (Fig. 6A). Note that the set of vertexes opposite to a given vertex in a *d*-simplex *κ*^(*d*)^ spans a (*d* – 1)-simplex—a *face* of *κ*^(*d*)^. The boundary of a *d*-simplex then consists of (*d* + 1) faces 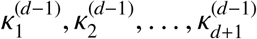, (Fig. 6B).
- *Geometric simplicial complexes* are combinations of geometric simplexes that match each other vertex-to-vertex, so that a non-empty intersection of any two simplexes in *K* yields another *K*-simplex: if *κ*_1_, *κ*_2_ ∈ *K*, then *κ*_1_ ∩ *κ*_2_ = *κ*_3_ ∈ *K*.
- The collection of all simplexes of dimensionality *d* and less forms the *d-skeleton* of *K*, *sk_d_*(*K*).
- Topological analyses of simplicial complexes do not address simplexes’ shapes and are based entirely on the combinatorics of the vertexes shared by the simplexes. This motivates using *abstract simplexes* and *abstract simplicial complexes* that capture the combinatorial structure of *κ*^(*d*)^s without making references to their geometry. Specifically, an abstract 0-simplex is a vertex 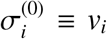, an abstract 1-simplex is a pair of vertexes, 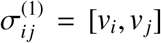; an abstract 2-simplex is a triple of vertexes, 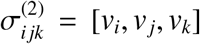, and so forth (Fig. 6C). Thus, abstract complexes may be viewed as multidimensional generalizations of graphs or as abstractions derived from the geometric simplicial complexes. A *d*-element subset of an abstract *d*-simplex *σ*^(*d*)^ forms its (*d* − 1)-face. The”face-matching” of the abstract simplexes in Σ means simply that a nonempty overlap of two simplexes *σ*_1_, *σ*_2_ ∈ Σ is a simplex of the same complex, *σ*_1_ ∩ *σ*_2_ = *σ*_3_ ∈ Σ. The latter property is commonly used to define abstract simplicial complexes for arbitrary sets, using families of their subsets that are closed under the “∩” operation [41].

- *Example 1*: The set of overlapping regions (4) define abstract simplexes (5) of the nerve complex (6) (Fig. 7A).
- *Example 2*: The combinations of coactive cells define coactivity simplexes (1), which together form a coactivity complex (Fig. 7B).
- *Example 3*. Vertexes of geometric simplexes that form a geometric simplicial complex *K* define abstract simplexes that form the corresponding abstract simplicial complex Σ (Fig. 7C).
- The set of *d*-dimensional simplexes of a complex Σ forms its (abstract) d-skeleton, *sk_d_*(Σ).
- *A clique complex* of an undirected graph *G* is an abstract simplicial complex formed by the cliques (fully interconnected subgraphs) of *G* [91], Fig. 6A. Combinatorial properties of cliques are the same as simplexes’: a subset of a clique’s vertexes form a clique, overlap of two *G*-cliques is also a clique, *ς*_1_ ∪ *ς*_2_ = *ς*_3_ ∈ *G* (Fig. 6). Thus, any graph *G* defines a unique clique complex 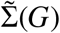. Note, that the 1-skeleton of a clique complex yields its underlying graph, 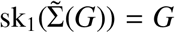, but if Σ is not a clique complex, then the clique complex built over its 1-skeleton does not reproduce Σ.
- *Coactivity complexes* used in this study are of two kinds. The first kind is formed by the abstract complexes 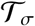 built from simultaneously coactive cell groups (1). The second kind is formed as the clique complexes of the coactivity graphs 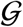 [77, 80]. The graph (co)activity thresholds *m*and μare used to control the size of the complex 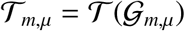: selecting *m* ≥ 2, *μ* = 1 for small maps (i.e., counting cells that produce at least two spikes per time bin *w*) and *m* ≥ 2, *μ* ≥ 5 for larger maps allows computing the full simplicial complex with dimensionality 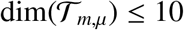, for which we can numerically apply the Javaplex software [119].

**FIG. 7:**
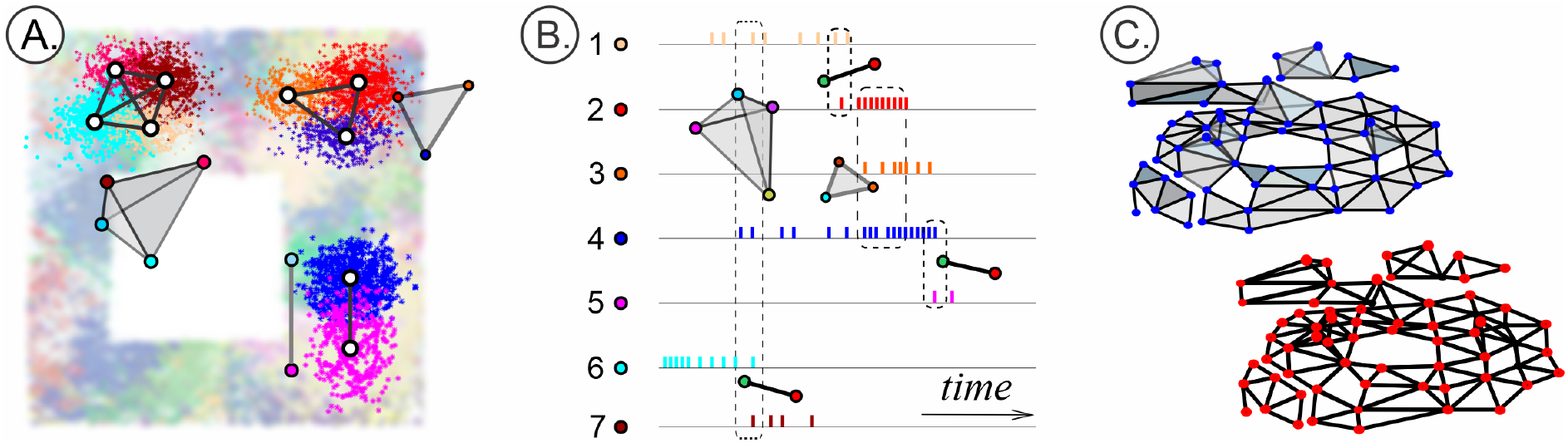
Cliques and simplexes. (**A**). Pairwise interlinked place fields produce cliques of the coactivity graph 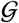. Shown is a vertex *ς*^(0)^ (0-clique), a link *ς*^(1)^ (1-clique), a three-vertex *ς*^(2)^ and a four-vertex *ς*^(3)^ clique. (**B**). Geometric simplexes: a 0D dot (*κ*^(0)^), a 1D link (*κ*^(1)^), a 2D triangle (*κ*^(2)^) and a 3D tetrahedron (*κ*^(3)^). (**C**). The corresponding complexes: a simplicial coactivity complex 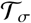 whose simplexes (1) are detected as singular coactivity events (left) may topologically differ from the clique coactivity complexes 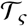, assembled from the cliques of a coactivity graph 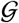 (right) over a spike integration period *ϖ*. A simplicial complex *K* is a combination of matching simplexes. The set of vertexes and black lines highlight the 1*D*-skeleton of *sk*_1_(*K*).

#### Topological invariants

- *Homological groups* are designed to “count pieces” in a space *X* with suitable coefficients. The key property of these groups is that they remain unchanged—*invariant*— as *X* is continuously deformed (see [41, 42] for a gentle introduction to the subject). If the coefficients form an algebraic field *F*, then the homological groups, commonly referred to as the “homologies” of *X* are simply vector spaces *H*_0_(*X*, *F*), *H*_1_(*X, F*),…, associated with *X* (one per dimension). Homologies can be easily computed for spaces whose “pieces” are explicitly defined, e.g., for the simplicial complexes, thus providing a way of identifying their topological structures. In practice, it is easier to use just the dimensionalities of *H*_*_s—the *Betti numbers b_k_* = dim(*H_k_*(*X, F*)), to count numbers of connectivity components, cavities, tunnels and other topological features of *X* in different dimensions [41, 42]. For example, if *X* is the boundary of a hollow triangle (or another noncontractible 1*D* loop), then *β*_1_(*X*) = 1. If *X* is 1-dimensional complex, i.e., a graph, then *β*_1_(*X*) equals to the number of cycles in *X*, counted up to topological equivalence. If the triangle is “filled”, then it can be continuously contracted into a 0*D* point; since the latter has no topological structure in dimensions *d* > 0, the corresponding Betti numbers also vanish. By the same argument a “filled” tetrahedron has *β*_*k*>0_ = 0, but if the tetrahedron is hollow, then its boundary, being a 2*D* noncontractible loop (topologically—a 2*D* sphere) produces *β*_2_ = 1, *β*_*k*>2_ = 0. Similarly, for any *d*-simplex *β*_*k*>0_(*σ*^(*d*)^) = 0, whereas for its hollow boundary, ∂*σ*^(*d*)^, the Betti numbers are *β*_*d*−1_(∂*σ*^(*d*)^) = 1, *β*_*k*≠0,*d*−1_(∂*σ*^(*d*)^) = 0 (Fig. 6). Same results apply to the “abstract” counterparts of all these complexes. Note also that continuous deformations of a 0*D* point *x* (a 0*D* topological loop) amount to “sliding”*x* inside of a space *X* that contains *x*; thus *β*_0_(*X*) simply counts such “sliding domains”, i.e., the number of connected components in *X*. As a result, all simplexes and simplicial complexes that consist of one piece have *β*_0_(*X*) = 1.
- *Persistent homology* theory allows tracing the topological structure in a filtered family of simplicial complexes, e.g., describing the topological dynamics of the inflating family (2), [70–73]. The Betti numbers plotted as function of the filtration parameter (in our case it is time, *t*) form the *barcode*, 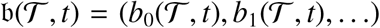, which provides the exact mathematical meaning to the term “topological shape” used throughout the text. Each bar in 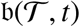 can be viewed as the corresponding topological loop’s timeline [48, 74–77, 80].
- *Zigzag Homology* theory allows tracking the Betti numbers of the “flickering” complexes— the ones whose simplexes can not only appear, but also disappear (see [110, 111] and Supplement in [107]). In particular, Zigzag homology techniques allow capturing the times when individual loops appear in the flickering complex, how long they persist, when they disappear, reappear again, etc.

#### Representability

A generic algorithm for checking whether a given complex can be built as a nerve of a *D*-dimensional cover is known only for *D* = 1 (see below). However, there exist criteria that allow ruling out certain non–representable cases.

- The *Leray criterion* posits that if a complex Σ is a nerve of a D-dimensional cover with contractible overlaps (4), then its rational homologies in dimensions higher or equal than *D* should vanish, 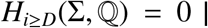 [58]. Moreover, homologies of all the subcomplexes Σ′ ⊆ Σ, induced by selecting vertex subsets of Σ should also vanish, 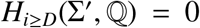. These properties can be verified by computing the Betti numbers and verifying that 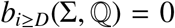. In practice, it is more convenient to carry out the computations over a finite field, such as 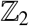. Although the 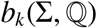 numbers may in general differ from the 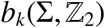 numbers, the latter also have to obey the Leray condition and produce the same Leray dimensionality. As an example, the Leray condition poses that the boundary of the triangle is not 1-representable 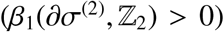, but the triangle itself may be 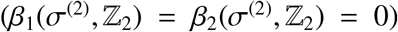; the boundary of a tetrahedron is not 2-representable 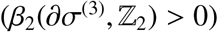, but the tetrahedron may be.
- *Amenta’s theorem* connects the Leray dimensionality of a simplicial complex to its *Helly number*, defined as follows. Let ϒ = {*ν*_1_, *ν*_2_, …, *ν_n_*} be a finite family of regions (Fig. 8). The Helly number *h* = *h*(ϒ) of the family is defined to be the maximal number of non-overlapping regions, such that every *h* − 1 among them overlap. For the corresponding nerve complex 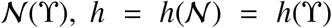 is the number of vertices of the largest simplicial hole in 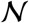 (i.e., the dimension of the hole plus 2, [49]). This observation can be used to attribute a Helly number to any simplicial complex Σ, *h*(Σ). From the perspective of representability analyses, a key property of the Helly numbers is that they do not exceed *d* + 1 for a *d*-Leray complex [49]. In particular, if the regions ν_*i*_ ∈ ϒ consist of up to *k* compact, convex domains in *R^d^*, and any intersection *β*_*i*_0__ ∩ … ∩ *ν*_i_t__ also satisfies this property, then h(ϒ) ≤*k*(*d*+ 1) [49, 105, 120].
- *Eckhoff ‘s conjecture*. The *f*-vector *f* = (*f*_1_, *f*_2_,…, *f_n_*) of a simplicial complex Σ is the list of numbers of its *k*-dimensional simplexes, *f_k_* = #{*σ_i_* ∈ Σ| dim(*σ*) = *k*}(“*f*” is a traditional notation that should not be confused with the firing rates). The *h*-vector of Σ is defined as

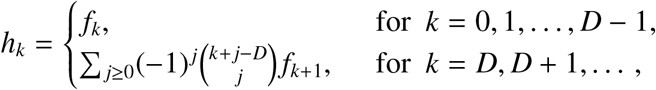

where parentheses denote the binomial coefficients. Given the combinatorial decomposition of *l*,

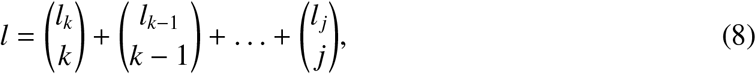

where *l_k_* ≥ *l*_*k*−1_ ≥ … ≥ *l_j_* ≥ *j* ≥ 1 [121], define the set of numbers

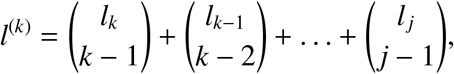

with 0^(*k*)^ = 0. Eckhoff’s conjecture [59], proven in [60] holds that the *h*-numbers of a *d*-representable complex must satisfy the following inequalities:

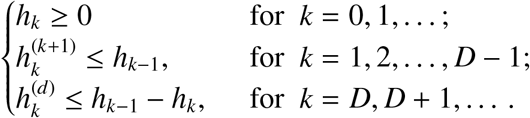

which can be verified not only for “static” complexes, but also for the “inflating” (2) and “flickering” complexes, at each step of their evolution.
- *Qualitative spatial consistency*. It can be shown that if the RCC5 relationships among all triples of regions are consistent, then the entire schema 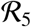 is consistent [93–98]. The full set of consistent triples is given in the following table.
- *Recognizing 1-representability* algorithm follows the exposition in [56, 57]. Let 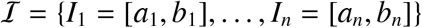 be set of intervals of a Euclidean line *R*^1^.

**FIG. 8:**
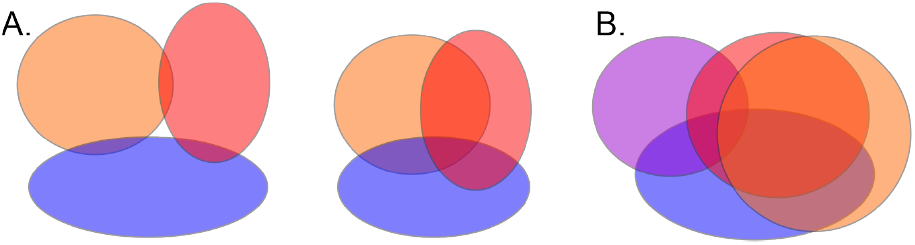
Helly’s theorem. (**A**). Three regions may exhibit both pairwise (left) or triple overlap (right). (**B**). The Helly number of a family of convex regions in 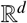 does not exceed *d* + 1. Thus, for convex planar regions having all triple overlaps implies having all the higher order (i.e., for this particular picture quadruple) overlaps. Hence, the intersection patterns of convex planar subspaces are completely determined by the intersection patterns of triples.

##### Definition 1.

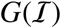 *is an interval graph, if each vertex 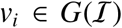 corresponds to an interval 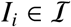 and a pair of vertexes* (*v_i_, v_j_*) *is connected by an edge iff I_i_ and I_j_ intersect*.

An interval graph is hence 1-dimensional skeleton of the nerve of 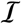 (Fig. 9A). It can also be verified that the complement of an interval graph is a directed comparability graph 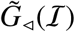, in which the relationship *v_i_* ◁ *v_j_* is defined by the order of the overlapping intervals,

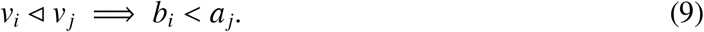

**FIG. 9:**
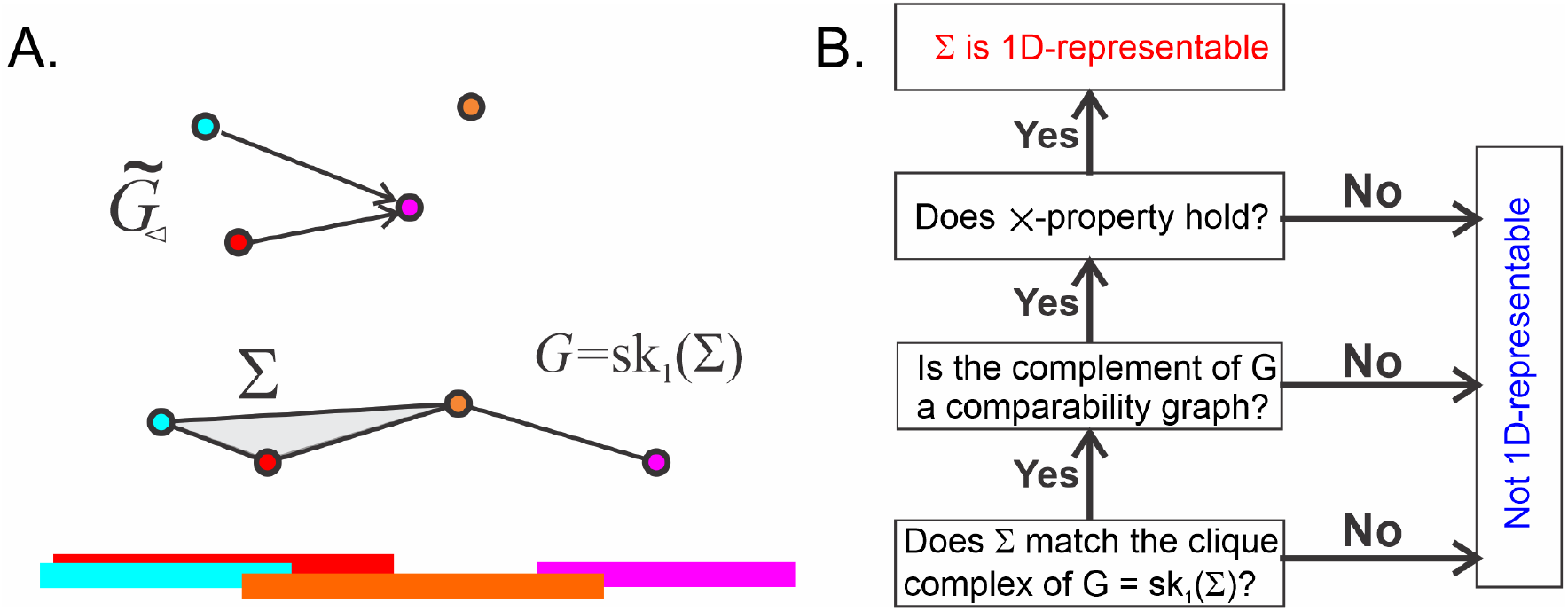
An algorithm for recognizing 1-representable complexes. **A**. Four intervals covering a linear segment (bottom) can be represented by a simplicial complex—the nerve of the cover (middle panel). The vertexes of the corresponding interval graph 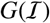—the 1*D* skeleton of Σ—(color-coded) are connected if their respective intervals overlap, *I_i_* ∩ *I_j_* = ⟹ *v_i_* ◁ *vj*. The corresponding comparability graph, 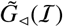 is shown above, with the order indicated by arrows: *v_i_* ◁ *v_j_* iff there is an arrow leading from *v_i_* to *v_j_*. **B**. Given a simplicial complex Σ, first check whether it is the clique complex of its 1*D*-skeleton *G* ≔ *sk*_1_(Σ). If it is not, then Σ is not 1-representable; if it is, then check whether the complement graph of *G* is a comparability graph. If it is not, then Σ is not 1–representable. If it is, then check the ×-property: if it holds, then Σ is 1–representable, otherwise it is not.

##### Definition 2.

*A directed graph satisfies Λ-property if there are no three vertices v_i_, v_j_, v_k_ such that v_i_, v_k_ are not adjacent, while v_i_ is adjacent to v_j_ and v_j_ is adjacent to v_k_ with the corresponding orientations being e_ij_ and e_jk_ respectively*.

##### Definition 3.

*An interval graph satisfies ×-property if no four vertexes v_i_, v_j_, v_k_, v_l_ produce disjoint pairs of intervals*.

In other words, a situation when *v_i_* ◁ *v_j_* and *v_k_* ◁ *v_l_* (i.e., the pair of intervals (*I_i_*, *I_j_*) overlaps and the pair (*I_k_*, *I_l_*) also overlaps), while the remaining pairs remain incompatible, e.g., *v_j_* ⋪ *v_k_*, *v_j_* ⋪ *v_l_*, (i.e., *I_j_* does not overlap either *I_k_* or *I_l_*), etc., does not appear.

##### Theorem.

*A graph is an interval graph iff its complement is a comparability graph with an order defined by (9), satisfying the ×-property*.

This theorem and the definitions motivate the following algorithm for identifying 1*D* representability of a complex Σ (Fig. 9B):

1. Test whether Σ is a clique complex, i.e., verify whether all (*k* + 1)-tuples of vertexes *v_σ_* = [*v*_*i*_0__, *v*_*i*_1__,…, *v_ik_*] form a simplex in Σ if and only if each pair of vertexes [*v_i_p__, v_i_q__*] ∈ *v_σ_* is an edge in its 1-skeleton *G* = *sk*_1_(Σ). If at least one *v_σ_* fails this test, then Σ is not a clique complex and hence not representable.
2. Build the complement 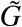 of *sk*_1_(Σ) and verify its comparability as follows:

i. Choose an edge between *v_i_* and *v_j_* and define an orientation on it (e.g., *e_ij_* ≠ *e_ji_*). If *e_i_j* was selected, then search for all vertexes *v*_*j*′_ that are connected to *V_j_* but not to *v_i_* (Fig. 9A). If the edge between *j* and *j*′ is not yet oriented, select *e*_*j*′*j*_. If it was already (*j*′ *j*)-oriented, continue on; the opposite, (*jj*′)-orientation implies that Σ is not representable. If the orientation for new edges cannot be selected, pause the algorithm and dispose of all the edges that have already been oriented. Then pick another unoriented edge and restart the Λ-rule: keep applying it until the process comes to a halt and the next set of edges needs to be removed. Do this until all the edges are serviced and hence oriented.
ii. Verify that no 3-tuple of vertexes (*v_i_*, *v_j_*, *v_k_*) forms an oriented 3-cycle. If such a cycle exists, Σ is non-representable in 1*D*.
iii. Verify that no triple of vertexes (*v_i_*, *v_j_*, *v_k_*) is “disconnected,” i.e., given *e_ij_* and *e_jk_*, there must exist an edge between *i* and *k*. If any triple violates this condition, Σ is not representable. Otherwise 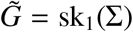 is a comparability graph with the order: *v_i_* ◁ *v_j_* for each *e_ij_*.
3. For every vertex *v_i_*, compute the set of lesser points, *D*(*v_i_*) = {*v_j_* ∈ *V*: *v_j_* ◁ *s_i_*}. Then, for all pairs of vertexes (v¿, *vj*) check whether D(v¿) is a subset of *D(vj*) or vice-versa. If at least one of these conditions is not satisfied, Σ is not representable.

If this sequence of conditions is satisfied, Σ is 1*D*-representable.

## Footnotes

1 Throughout the text, terminological definitions are given in *italics*.

## REFERENCES

[1] O’Keefe, J. & Dostrovsky, J. The hippocampus as a spatial map. Preliminary evidence from unit activity in the freely-moving rat. Brain Res. 34(1): 171–5 (1971).

[2] O’Keefe, J. Place units in the hippocampus of the freely moving rat. Experimental Neurology. 51(1): 78–109 (1976).

[3] Vinogradova, O. Functional Organization of the Limbic System in the Process of Registration of Information: Facts and Hypotheses. In: Isaacson RL, Pribram KH, editors. The Hippocampus 2: 3–69. Springer, Boston (1975).

[4] Best, P., White, A., Minai, A. (2001). Spatial processing in the brain: the activity of hippocampal place cells. Annual. Rev. Neurosci. 24: 459–486.

[5] Brown, E., Frank, L., Tang, D., Quirk, M. & Wilson, M. A statistical paradigm for neural spike train decoding applied to position prediction from ensemble firing patterns of rat hippocampal place cells. J. Neurosci., 18: 7411–7425 (1998).

[6] Barbieri, R., Wilson, M.A., Frank, L.M. & Brown, E.N. An analysis of hippocampal spatio-temporal representations using a Bayesian algorithm for neural spike train decoding. IEEE transactions on neural systems and rehabilitation engineering 13: 131–136 (2005).

[7] Jensen, O. & Lisman, J.E. Position reconstruction from an ensemble of hippocampal place cells: contribution of theta phase coding. J. Neurophysiol. 83: 2602–2609 (2000).

[8] Guger, C., Gener, T., Pennartz, C., Brotons-Mas, J., Edlinger, G., Bermúdez, I., Badia, S., Verschure, P., Schaffelhofer, S. & Sanchez-Vives MV. Real-time position reconstruction with hippocampal place cells. Front. Neurosci., 5: 85 (2011).

[9] Karlsson, M. & Frank L. Awake replay of remote experiences in the hippocampus. Nat. Neurosci. 12: 913–918 (2009).

[10] Wu, X. & Foster, D. Hippocampal Replay Captures the Unique Topological Structure of a Novel Environment. J. Neurosci. 34: 6459–6469 (2014).

[11] Ji, D. & Wilson, M. Coordinated memory replay in the visual cortex and hippocampus during sleep. Nat. Neurosci. 10: 100–107 (2007).

[12] Johnson, A. & Redish, A. Neural Ensembles in CA3 Transiently Encode Paths Forward of the Animal at a Decision Point. J. Neurosci., 27, 12176–12189 (2007).

[13] Dragoi, G. & Tonegawa, S. Preplay of future place cell sequences by hippocampal cellular assemblies. Nature 469: 397–401 (2011).

[14] Pfeiffer, B. & Foster, D. Hippocampal place-cell sequences depict future paths to remembered goals. Nature 497: 74–9 (2013).

[15] Moser, E., Moser, M-B. & McNaughton, B. Spatial representation in the hippocampal formation: a history. Nat. Neurosci. 20(11): 1448–64 (2017).

[16] Derdikman, D. & Moser, E. A manifold of spatial maps in the brain. Trends in Cognitive Sciences 14(12): 561–9 (2010).

[17] Grieves, R. & Jeffery, K. The representation of space in the brain. Behavioural Processes. 135: 113–31 (2017).

[18] Moser, E., Kropff, E. & Moser M-B. Place Cells, Grid Cells, and the Brain’s Spatial Representation System. Annu Rev. Neurosci. 31(1): 69–89 (2008).

[19] Taube J., Muller R., Ranck J., Jr. Head-direction cells recorded from the postsubiculum in freely moving rats. J. Neurosci. 10: 420–435, 436–447 (1990).

[20] Taube, J., Goodridge, J., Golob, E., Dudchenko, P. & Stackman, R. Processing the head direction cell signal: a review and commentary. Brain Res. Bull. 40: 477–484 (1996).

[21] Wiener, S., & Taube, J. (Eds.). Head direction cells and the neural mechanisms of spatial orientation. MIT Press (2005).

[22] Savelli, F. & Knierim, J. Origin and role of path integration in the cognitive representations of the hippocampus: Computational insights into open questions. J. Exp. Biology, 222:jeb188912 (2019).

[23] Jeffery, K., Wilson, J., Casali, G. & Hayman, R. Neural encoding of large-scale three-dimensional space-properties and constraints. Front Psychol. 6: 927–939 (2015).

[24] Knierim, J. & McNaughton, B. Hippocampal place-cell firing during movement in three-dimensional space. J Neurophysiol. 85(1): 105–16 (2001).

[25] Hayman, R., Verriotis, M., Jovalekic, A., Fenton, A. & Jeffery, K. Anisotropic encoding of threedimensional space by place cells and grid cells. Nat Neurosci. 14(9): 1182–8 (2011).

[26] Grieves, R., Jedidi-Ayoub, S., Mishchanchuk, K., Liu, A., Renaudineau, S. & Jeffery, K. The placecell representation of volumetric space in rats. Nat Commun. 11(1):789 (2020).

[27] Rubin. A., Yartsev M., & Ulanovsky, N. Encoding of Head Direction by Hippocampal Place Cells in Bats. J. Neurosci. 34: 1067–1080 (2014).

[28] Finkelstein, A., Derdikman, D., Rubin, A., Foerster, J., Las, L. & Ulanovsky, N. Three-dimensional head-direction coding in the bat brain. Nature. 517(7533): 159–64 (2015).

[29] Georges-François, P., Rolls, E. & Robertson, R. Spatial View Cells in the Primate Hippocampus: Allocentric View not Head Direction or Eye Position or Place. Cerebral Cortex, 9(3): 197–212 (1999).

[30] Rolls, E., Robertson, R., Georges-François, P. Spatial View Cells in the Primate Hippocampus. European Journal of Neuroscience, 9(8): 1789–94 (1997). https://doi.org/10.1016/j.bbr.2010.03.027.

[31] Rolls, E. Spatial view cells and the representation of place in the primate hippocampus. Hippocampus 9(4): 467–80 (1999).

[32] Buffalo, E. Bridging the Gap Between Spatial and Mnemonic Views of the Hippocampal Formation. Hippocampus, 25(6): 713–8 (2015).

[33] de Araujo, I., Rolls, E. & Stringer, S. A view model which accounts for the spatial fields of hippocampal primate spatial view cells and rat place cells. Hippocampus 11(6): 699–706 (2001).

[34] Hubel, D. & Wiesel, T. Receptive fields of single neurones in the cat’s striate cortex. J. Physiology 148(3): 574–91 (1959).

[35] Arun, P., Sripati, A., Yoshioka, T., Denchev, P., Hsiao, S. & Johnson, K. Spatiotemporal Receptive Fields of Peripheral Afferents and Cortical Area 3b and 1 Neurons in the Primate Somatosensory System, J. Neurosci. 26: 2101–2114 (2006),

[36] Aertsen, A. & Johannesma, P. The Spectro-Temporal Receptive Field. Biol. Cybern. 42: 133–143 (1981).

[37] Atencio, C., Sharpee, T., Schreiner, C. Cooperative Nonlinearities in Auditory Cortical Neurons. Neuron. 58(6): 956–66 (2008).

[38] Gosselin, F. & Schyns, P. RAP: a new framework for visual categorization. Trends in Cognitive Sciences. 6(2): 70–7 (2002).

[39] DeAngelis, G, Ohzawa, I., Freeman, R. Receptive-field dynamics in the central visual pathways. Trends in Neurosciences. 18(10): 451–8 (1995).

[40] Babichev, A., Cheng, S. & Dabaghian, Y. Topological schemas of cognitive maps and spatial learning. Front. Comput. Neurosci. 10: 18 (2016).

[41] Aleksandrov, P. Elementary concepts of topology. (F. Ungar Publishing, 1965).

[42] Hatcher, A. Algebraic topology. Cambridge; New York: Cambridge University Press (2002).

[43] Alexandroff, P. Untersuchungen über Gestalt und Lage abgeschlossener Mengen beliebiger Dimension. Annals of Mathematics., 30, 101–187 (1928).

[44] Čech, E. Théorie générale de l’homologie dans un espace quelconque. Fundamenta mathematicae, 19, 149–183 (1932).

[45] Edwards, D. & Hastings, H. Čech Theory: Its past, present, and future. Rocky Mountain J. Math. 10(3): 429–468 (1980).

[46] De Silva, V. & Ghrist, R. Coverage in sensor networks via persistent homology. Algebraic & Geometric Topology 7: 339–358 (2007).

[47] Curto, C. & Itskov, V. Cell groups reveal structure of stimulus space, PLoS Comput. Biol., 4: e1000205 (2008).

[48] Dabaghian, Y., MéEmoli, F., Frank, L. & Carlsson, G. A Topological Paradigm for Hippocampal Spatial Map Formation Using Persistent Homology, PLoS Comput. Biol., 8: e1002581 (2012).

[49] Tancer, M. Intersection Patterns of Convex Sets via Simplicial Complexes: A Survey. In: Pach J, Ed. Thirty Essays on Geometric Graph Theory: Springer New York. pp. 521–40 (2013).

[50] Tancer, M. d-Representability of simplicial complexes of fixed dimension. Journal of Computational Geometry. 2(1): 183–8 (2011).

[51] Kratochvíl, J. & Matoušek, J. Intersection graphs of segments. J. Comb. Theory Ser. B, 62(2): 289–315 (1994).

[52] Matousek, J, Tancer, M. & Wagner, U. Hardness of embedding simplicial complexes in *R^d^*. Proceedings of the twentieth Annual ACM-SIAM Symposium on Discrete Algorithms; New York, New York. 1496863: Society for Industrial and Applied Mathematics, p. 855–64 (2009).

[53] Fulkerson, D. & Gross, O. Incidence matrices and interval graphs. Pacific J. Math. 15(3): 835–855 (1965).

[54] Habib, M., McConnell, R., Paul, C. & Viennot, L. Lex-BFS and partition refinement, with applications to transitive orientation, interval graph recognition and consecutive ones testing. Theoretical Computer Science. 234(1): 59–84 (2000).

[55] Kratsch, D., McConnell, R., Mehlhorn, K. & Spinrad, J. Certifying Algorithms for Recognizing Interval Graphs and Permutation Graphs. SIAM Journal on Computing 36(2): 326–353 (2006).

[56] Golumbic, M. The Complexity of Comparability Graph Recognition and Coloring. Computing 18: 199–208 (1977).

[57] Fishburn, P. Interval graphs and interval orders. Discrete Mathematics 55: 135–149 (1985).

[58] Leray, J. Sur la forme des espaces topologiques et sur les points fixes des représentations. J. Math. Pures Appl, 24: 95–167 (1945).

[59] J. Eckhoff, Über kombinatorisch-geometrische Eigenschaften von Komplexen and Familien knovexer Mengen, J. Reine Angew. Math, 313: 171–188 (1980).

[60] Kalai, G. Characterization of *f*-vectors of families of convex sets in *R^d^* part II: Sufficiency of Eckhoff ‘s conditions. Journal of Combinatorial Theory, Series A. 41(2): 167–88 (1986).

[61] Kalai, G. Intersection patterns of convex sets. Israel Journal of Mathematics, 48(2-3): 161–74 (1984).

[62] Kalai, G. & R. Meshulam. Leray numbers of projections and a topological Helly type theorem. J. Topology, 1(3): 551–556 (2008).

[63] Kalai, G. & Meshulam, R. A topological colorful Helly theorem. Adv. Math., 191(2): 305–311 (2005).

[64] Kalai, G. & Meshulam, R. Intersections of Leray complexes and regularity of monomial ideals. J. Combin. Theory Ser. A, 113(7): 1586–1592 (2006).

[65] Tancer, M. & Tonkonog, D. Nerves of Good Covers Are Algorithmically Unrecognizable. SIAM Journal on Computing, 42(4): 1697–719 (2013).

[66] T. Hafting, M. Fyhn, S. Molden, M.-B. Moser and E.I. Moser. Microstructure of a spatial map in the entorhinal cortex. Nature 436: 801–806 (2005).

[67] Brun, V., Solstad, T., Kjelstrup, K., Fyhn, M., Witter, M., Moser, E. & Moser, M-B. Progressive increase in grid scale from dorsal to ventral medial entorhinal cortex.

[68] Ghrist, R. Barcodes: The persistent topology of data, Bull. Amer. Math. Soc., 45: 61–75 (2008).

[69] Kang, L., Xu. B. & Morozov, D. Evaluating State Space Discovery by Persistent Cohomology in the Spatial Representation System. Front. Comput. Neurosci. 15(28):616748 (2021).

[70] Wasserman, L. Topological Data Analysis. Annual Review of Statistics and Its Application 5: 501–532 (2018).

[71] Zomorodian, A., & Carlsson, G. Computing persistent homology. Discrete Comput Geom 33: 249–274 (2005).

[72] Edelsbrunner, H., Letscher, D., & Zomorodian, A. Topological Persistence and Simplification. Discrete & Computational Geometry 28: 511–533 (2002).

[73] Zomorodian, A. Topology for Computing Cambridge University Press, New York (2009).

[74] Arai, M., Brandt, V. & Dabaghian Y. The Effects of Theta Precession on Spatial Learning and Simplicial Complex Dynamics in a Topological Model of the Hippocampal Spatial Map. PLoS Comput Biol. 10: e1003651 (2014).

[75] Basso, E., Arai, M. & Dabaghian, Y. The effects of gamma synchronization on spatial learning in a topological model of the hippocampal spatial map. PloS Comput. Biol. 12: 9 (2016).

[76] Dabaghian, Y. Through synapses to spatial memory maps: a topological model. Sci. Reports 9: 572 (2018).

[77] Babichev, A., Mémoli, F., Ji, D. & Dabaghian, Y. A topological model of the hippocampal cell simplex network. Frontiers in Comput. Neurosci., 10: 50 (2016).

[78] Babichev, A. & Dabaghian, Y. Transient cell simplex networks encode stable spatial memories. Sci. Rep. 7: 3959 (2017).

[79] Dabaghian, Y. From Topological Analyses to Functional Modeling: The Case of Hippocampus. Front. Comput. Neurosci. 14 (2021).

[80] Hoffman, K., Babichev, A. & Dabaghian, Y. A model of topological mapping of space in bat hippocampus. Hippocampus, 26: 1345–1353 (2016).

[81] Lisman, J. Buzsáki, G., Eichenbaum, H., Nadel, L., Ranganath, C. & Redish, A. Viewpoints: how the hippocampus contributes to memory, navigation and cognition. Nature Neuroscience. 20: 1434–48 (2017).

[82] Y. Dabaghian, Learning Orientations: a Discrete Geometry Model, in submission.

[83] Curto, C. & Vera, R. The Leray Dimension of a Convex Code. arXiv:1612.07797 (2016).

[84] Buzsáki, G. Neural syntax: cell assemblies, synapsembles, and readers. Neuron 68: 362–385 (2010).

[85] König, P., Engel, A. & Singer, W. Integrator or coincidence detector? The role of the cortical neuron revisited. Trends Neurosci., 19: 130–137 (1996).

[86] London, M. & Häusser, M. Dendritic Computation. Ann. Rev. Neurosci. 28: 503–532 (2005).

[87] Koulakov, A.A., Raghavachari, S., Kepecs, A. & Lisman, J.E. Model for a robust neural integrator. Nat Neurosci 5, 775–782 (2002).

[88] Spruston, N. Pyramidal neurons: dendritic structure and synaptic integration. Nat. Rev. Neurosci. 9: 206–221 (2008).

[89] Burgess, N. & O’Keefe, J. Cognitive graphs, resistive grids, and the hippocampal representation of space. J. Gen. Physiol. 107: 659–662 (1996).

[90] Muller, R., Stead, M. & Pach, J. The hippocampus as a cognitive graph. J. Gen. Physiol. 107: 663–694 (1996).

[91] Jonsson, J. Simplicial complexes of graphs, Springer, New York (2008).

[92] Mizuseki, K., Sirota, A., Pastalkova, E. & Buzsáki, G. Theta oscillations provide temporal windows for local circuit computation in the entorhinal-hippocampal loop, Neuron, 64: 267–280 (2009).

[93] Cohn, A.G. & Renz., J. Qualitative Spatial Representation and Reasoning, in Foundations of Artificial Intelligence, van Harmelen, F., Lifschitz, V. & Porter, B. (Eds), Elsevier. pp. 551–596 (2008).

[94] Chen, J., Cohn, A., Liu, D., Wang, S., Ouyang, J., & Yu, Q. A survey of qualitative spatial representations. Knowledge Engineering Rev., 30(1), 106–136 (2015).

[95] A. G. Cohn and N. M. Gotts, Spatial Regions with Undetermined Boundaries, Proceedings of Gaithesburg Workshop on GIS, ACM (1994).

[96] J. Renz, A Canonical Model of the Region Connection Calculus, Journal of Applied Non-Classical Logics, 12(3-4): 469–494, (2002)

[97] B. Bennett, Determining Consistency of Topological Relations, Constraints 3(2-3): 213–225, (1998).

[98] Long Z. & Li S. On Distributive Subalgebras of Qualitative Spatial and Temporal Calculi. In: Fabrikant S., Raubal M., Bertolotto M., Davies C., Freundschuh S., Bell S. (eds) Spatial Information Theory. COSIT 2015. Lecture Notes in Computer Science, vol 9368. Springer, Cham (2015).

[99] Brown, E., Nguyen, D., Frank, L., Wilson, M. & Solo, V. An analysis of neural receptive field plasticity by point process adaptive filtering. Proc. Natl Acad. Sci. 98: 12261–66 (2001).

[100] Barbieri, R., Frank, L., Nguyen, D., Quirk, M., Solo V, et al. Dynamic analyses of information encoding in neural ensembles. Neural Comput. 16: 277–307 (2004).

[101] U. Eden, L. Frank, R. Barbieri, V. Solo and E. Brown, Dynamic analysis of neural encoding by point process adaptive filtering Neural Comput 16: 971–998 (2004).

[102] Frank, L, Brown, E & Stanley, G. Hippocampal and cortical place cell plasticity: implications for episodic memory. Hippocampus 16: 775–784 (2006).

[103] Singer, A., Karlsson, M., Nathe, A., Carr, M. & Frank, L. Experience-dependent development of coordinated hippocampal spatial activity representing the similarity of related locations. J Neurosci. 30: 11586–11604 (2010).

[104] Knierim, J. Dynamic Interactions between Local Surface Cues, Distal Landmarks, and Intrinsic Circuitry in Hippocampal Place Cells. J. Neurosci. 22: 6254–6264 (2002).

[105] Amenta, N. A short proof of an interesting Helly-type theorem. Discrete Comput. Geom, 15: 423–427, (1996).

[106] Curto, C. et al. What Makes a Neural Code Convex? SIAM Journal on Applied Algebra and Geometry 1: 222–238 (2017).

[107] Babichev, A., Morozov, D. & Dabaghian, Y. Robust spatial memory maps encoded by networks with transient connections. PLoS Comput. Bio. 14(9): e1006433 (2018).

[108] Babichev, A., Morozov, D. & Dabaghian, Y. Replays of spatial memories suppress topological fluctuations in cognitive map. Network Neuroscience, Special Issue: Topological Neuroscience, 3(3): 707–724 (2019).

[109] Fenton A, Muller R. Place cell discharge is extremely variable during individual passes of the rat through the firing field. Proc. Natl. Acad. Sci. 95(6): 3182–3187 (1998).

[110] Carlsson, G. & Silva, Vd. Zigzag Persistence. Found. Comput. Math. 10: 367–405 (2010).

[111] Carlsson, G., Silva, Vd. & Morozov, D. Zigzag persistent homology and real-valued functions. Proceedings of the 25th annual symposium on Computational geometry. Aarhus, Denmark: ACM. pp. 247–256 (2009).

[112] Dabaghian, Y., Brandt, V. & Frank, L. Reconceiving the hippocampal map as a topological template, eLife 10.7554/eLife.03476: 1–17 (2014).

[113] Dabaghian, Y., Cohn, A. & Frank, L. Topological maps from signals. in Proceedings of the 15th ACM International Symposium on Geographic Information Systems, ACM-GIS 2007, November 7-9, Seattle, WA (61): 61–67 (2007).

[114] Battaglia, F, Sutherland, G. & McNaughton, B. Local sensory cues and place cell directionality: additional evidence of prospective coding in the hippocampus. J. Neurosci. 24: 4541–4550 (2004).

[115] Chazal F. & Yann Oudot, S. Towards persistence-based reconstruction in Euclidean spaces. In Proceedings of the Twenty-fourth Annual Symposium on Computational Geometry, SCG ‘08: 232–241, New York (2008).

[116] Cavanna, N. & Sheehy, D. The Generalized Persistent Nerve Theorem, arXiv:1807.07920.

[117] Eichenbaum, H., Dudchenko, P., Wood, E., Shapiro, M. & Tanila, H. The hippocampus, memory, and place cells: is it spatial memory or a memory space? Neuron 23: 209–226 (1999).

[118] Buzsáki, G. & Mizuseki, K. The log-dynamic brain: how skewed distributions affect network operations. Nat Rev Neurosci. 15(4): 264–278 (2014).

[119] Adams, H., Tausz, A., Vejdemo-Johansson, M. javaPlex: A Research Software Package for Persistent (Co)Homology. In: Hong H., Yap C. (eds) Mathematical Software – ICMS 2014. ICMS 2014. Lecture Notes in Computer Science, vol 8592. Springer, Berlin, Heidelberg. (2014)

[120] Danzer, L., Grünbaum, B. & Klee, V. Helly’s theorem and its relatives. Proc. Symp. Pure Math., 7: 101–180 (1963).

[121] Beckenbach, E. (Ed.) Applied Combinatorial Mathematics, pp. 27–30 (1964).

